# Structures of plasmepsin X from *P. falciparum* reveal a novel inactivation mechanism of the zymogen and molecular basis for binding of inhibitors in mature enzyme

**DOI:** 10.1101/2021.09.24.460453

**Authors:** Pooja Kesari, Anuradha Deshmukh, Nikhil Pahelkar, Abhishek B. Suryawanshi, Ishan Rathore, Vandana Mishra, John H. Dupuis, Huogen Xiao, Alla Gustchina, Jan Abendroth, Mehdi Labaied, Rickey Y. Yada, Alexander Wlodawer, Thomas E. Edwards, Donald D. Lorimer, Prasenjit Bhaumik

**Author notes:** Corresponding authors: Donald D Lorimer, Prasenjit Bhaumik. These authors contributed equally (Pooja Kesari, Anuradha Deshmukh).

## Abstract

*Plasmodium falciparum* plasmepsin X (*Pf*PMX), involved in the invasion and egress of this deadliest malarial parasite, is essential for its survival and hence considered as an important drug target. We report the first crystal structure of *Pf*PMX zymogen containing a novel fold of its prosegment. A unique twisted loop from the prosegment and arginine 244 from the mature enzyme are involved in zymogen inactivation; such mechanism, not previously reported, might be common for apicomplexan proteases similar to *Pf*PMX. The maturation of *Pf*PMX zymogen occurs through cleavage of its prosegment at multiple sites. Our data provide thorough insights into the mode of binding of a substrate and a potent inhibitor 49c to *Pf*PMX. We present molecular details of inactivation, maturation, and inhibition of *Pf*PMX that should aid in the development of potent inhibitors against pepsin-like aspartic proteases from apicomplexan parasites.

## Main

The phylum Apicomplexa comprises unicellular parasitic protists capable of infecting different animals and mammals. Among these parasites, *Plasmodium, Toxoplasma*, and *Cryptosporidium* are of major global health concern as they infect humans and animals^1,2^. *Plasmodium falciparum* is the most lethal species in the genus *Plasmodium*, responsible for human malaria, one of the deadliest diseases worldwide, causing deaths of thousands of people every year^3^. *Toxoplasma gondii* is an obligate intracellular parasite that causes a fatal disease toxoplasmosis^4,5^. *Cryptosporidium* species cause cryptosporidiosis, the second most prevalent form of infant diarrheal disease^6^. Despite the high incidence of these diseases, treatment options are limited due to low efficiency of the available drugs and continual emergence of the resistant strains. For such reasons there is a pressing need for novel, long-lasting therapeutic agents against different apicomplexan parasites^7,8,9^.

Pepsin-like aspartic proteases are involved in nutrient uptake, immune evasion, invasion, and egress, which are vital processes that enable parasites to cause successful infection inside the host cell. *P. falciparum* encodes ten pepsin-like aspartic proteases known as plasmepsins (PMs), *Pf*PMI to *Pf*PMX. Among them, *Pf*PMI, *Pf*PMII, *Pf*PMIV, and histo-aspartic protease (*Pf*PMIII or HAP), also referred to as vacuolar plasmepsins, are responsible for the degradation of host hemoglobin^10^. *Pf*PMV acts as a maturase for proteins exported into the red blood cells (RBC) cytoplasm, involved in host cell remodeling^11^. *Pf*PMIX and *Pf*PMX are involved in the invasion and egress of the parasite^12,13^. *T. gondii* encodes seven pepsin-like aspartic proteases called toxomepsins (*Tg*ASP1-*Tg*ASP7); of which *Tg*ASP3 shares 40% homology with *Pf*PMX and carries out a similar function. Invasion and egress, facilitated by proteases like *Tg*ASP3 and *Pf*PMX, are two key events essential for survival and dissemination of infection for these classes of parasites; therefore, these enzymes (later referred to as PMX-like proteases) are attractive drug targets^14,15,16,17,18^.

Mature *Pf*PMX acts on exoneme and microneme proteins such as SUB1, AMA1, reticulocyte-binding protein homologs (Rhs), and erythrocyte binding-like (EBL) proteins that are responsible for both invasion and egress. The Rh2N peptide has been reported to have faster cleavage rate as compared to other known substrates for this enzyme^19^. *Pf*PMX is expressed in the blood-stage schizont and merozoite stage; additionally, it is found in the gametocyte and liver stage of infection^12,13^. *Tg*ASP3 acts on microneme and rhoptry proteins^15^. Furthermore, *Pf*PMX and other similar proteases (PMX-like) from apicomplexan parasites share significant functional and sequence level similarity, hence understanding the molecular basis of the functional properties of these enzymes would undoubtedly be useful in the development of potent therapeutic agents against these deadly parasitic diseases.

The complete polypeptide of *Pf*PMX is synthesized and folded as inactive zymogen (*Pf*pro*-* PMX) consisting of a prosegment and a mature domain (*Pf*m-PMX). The prosegment of *Pf*PMX is longer than its counterpart in vacuolar plasmepsins and other gastric or plant pepsin-like aspartic proteases^12,20,21^. In general, activation of the zymogens of pepsin-like aspartic proteases takes place by removal of the prosegment under acidic conditions. It is suggested that vacuolar PMs have an auto-activation mechanism wherein the dimeric form is converted to a monomer under acidic conditions due to the disruption of the Tyr-Asp loop at the pro-mature junction, with subsequent unfolding and cleavage of the prosegment by the trans-activation process^22^. Conversely, in other pepsin-like aspartic proteases, the interaction of a lysine residue with the catalytic aspartates is responsible for inactivation of the zymogen, which then detaches from the active site under acidic pH, leading to the unfolding of prosegment followed by cleavage by the same enzyme molecule^20,23^. It has been shown that *Pf*ERC (*Plasmodium falciparum* endoplasmic reticulum-resident calcium-binding protein) protein works at the upstream position in the activation pathway of *Pf*PMX^24^. Upon activation, the zymogens of PMX-like proteases form mature enzymes which promote invasion and egress of the parasites. The longer length of the prosegment of *Pf*PMX zymogen indicates a possibility of a novel inactivation mechanism.

Inhibition studies with hydroxyethyl amine compound 49c, a peptidomimetic molecule found to inhibit *Pf*PMIX, *Pf*PMX, and *Tg*ASP3, and ultimately the invasion and egress of the parasites by impairing maturation of protein cascade, underscore the importance of these enzymes in parasite survival^12,15,18^. A previous study showed the importance of a phenylalanine residue from the flap in binding the inhibitor 49c^18^. However, detailed analysis of the interactions responsible for binding of substrate and inhibitors in the active site of these enzymes is still lacking. Interestingly, pepstatin A, which is a potent inhibitor of pepsin-like aspartic proteases, does not inhibit *Pf*PMIX and *Pf*PMX^12^; hence such observation demands further understanding of the molecular insights into the active sites of these enzymes. No experimental structure for these enzymes has been available so far. Therefore, structural and biochemical studies are needed to investigate the active site architecture, specificity and affinity of the substrates, as well as the inhibitors. Such studies should help in the development of potent lead compounds.

The present study reports the first crystal structure of *P. falciparum Pf*PMX zymogen (*Pf*pro-PMX) with a truncated prosegment. The prosegment of *Pf*pro-PMX is folded in a unique way, not observed in any other pepsin-like aspartic proteases. The zymogen exhibits a novel mechanism of inactivation among plasmepsins. Molecular dynamics, as well as structural analysis, elucidate for the first time the detailed molecular basis of substrate binding to *P. falciparum* mature PMX *(Pf*m-PMX) and its inhibition by a potent inhibitor 49c. Molecular details presented in this study should aid in the development of potent inhibitors against PMX-like proteases, with the aim to combat diseases caused by apicomplexan parasites.

## Results

### Crystal structure of *Pf*PMX zymogen (*Pf*pro-PMX)

A polypeptide (**Fig. 1a**) of *Pf*PMX, with truncated prosegment (R28p-N234p, where ‘p’ indicates the residues from the prosegment) and TEV cleavage site (ENLYFQ) at the C terminus was crystallized. From here onwards the zymogen obtained from our construct 1 (R28p-N573) is referred to as *Pf*pro-PMX and the mature enzyme activated or modeled from *Pf*pro-PMX is referred to as *Pf*m-PMX (I235-N573). Crystal structure of *Pf*pro-PMX has been determined at 2.1 Å resolution (**Table 1**). **Figure 1b** shows the sigma-A weighted 2*F*_*O*_*-F*_*C*_ electron density of the prosegment (**see Supplementary movie 1**). The first residue (I235) of *Pf*m-PMX is defined based on its sequence alignment with pepsinogen (**Extended Data Fig. 1a**). The polypeptide regions with residues K69p-N70p and E132p-K224p in the prosegment and the C-terminal linker region could not be modeled due to lack of features in the electron density. With the exception of a few residues (F308-S315) in the flap, E343-D352 of loop N2, and the last residue N573, the complete mature enzyme portion of the *Pf*pro-PMX structure could be modeled and refined. An N-acetylglucosamine molecule is visible in the well-defined electron density (**Supplementary Fig.1**), indicating that the protein was glycosylated during its expression.

**Table 1:** Data collection and refinement statistics of *Pf*pro-PMX structure.

| **Data collection statistics<sup>a</sup>** | ***Pfpro*-PMX** |
| --- | --- |
| Space group | *C* 2 |
| Unit cell parameters<br>a, b, c (Å)<br>$\alpha$, $\beta$, $\gamma$ (°) | 192.30, 50.88, 49.92<br>90, 93.15, 90 |
| Temperature (K) | 100 |
| Wavelength (Å) | 0.97872 |
| Resolution (Å) | 40-2.10 (2.20-2.10) |
| R<sub>merge</sub> (%) | 4.6 (120.4) |
| Completeness (%) | 99.8 (99.6) |
| Mean I/$\sigma$(I) | 15.4 (1.1) |
| Total reflections | 112,116 (7462) |
| Unique reflections | 28,381 (2079) |
| Redundancy | 4.0 (3.6) |
| B factor from Wilson plot (Å<sup>2</sup>) | 32.6 |
| No. of molecules in the asymmetric unit | 1 |
| **Refinement statistics** |  |
| Resolution (Å) | 35-2.1 |
| Working set: no. of reflections | 28299 |
| R<sub>factor</sub> (%) | 20.6 |
| Test set: no. of reflections | 2120 |
| R<sub>free</sub> (%) | 23.8 |
| Protein atoms | 3225 |
| **Geometry statistics** |  |
| R.M.S.D. bond distance (Å) | 0.007 |
| R.M.S.D. bond angle (°) | 0.800 |
| Isotropic average B-factor (Å<sup>2</sup>) |  |
| Protein | 70.14 |
| Ramachandran plot (%)<sup>*</sup> |  |
| Allowed regions | 96.22 |
| Generously allowed regions | 3.78 |
| Disallowed regions | 0 |
| PDB ID | 7RY7 |

<sup>a</sup>Values in parentheses correspond the highest resolution shell.
<sup>\*</sup>Ramachandran plot statistics based on MolProbity

**Figure 1:**
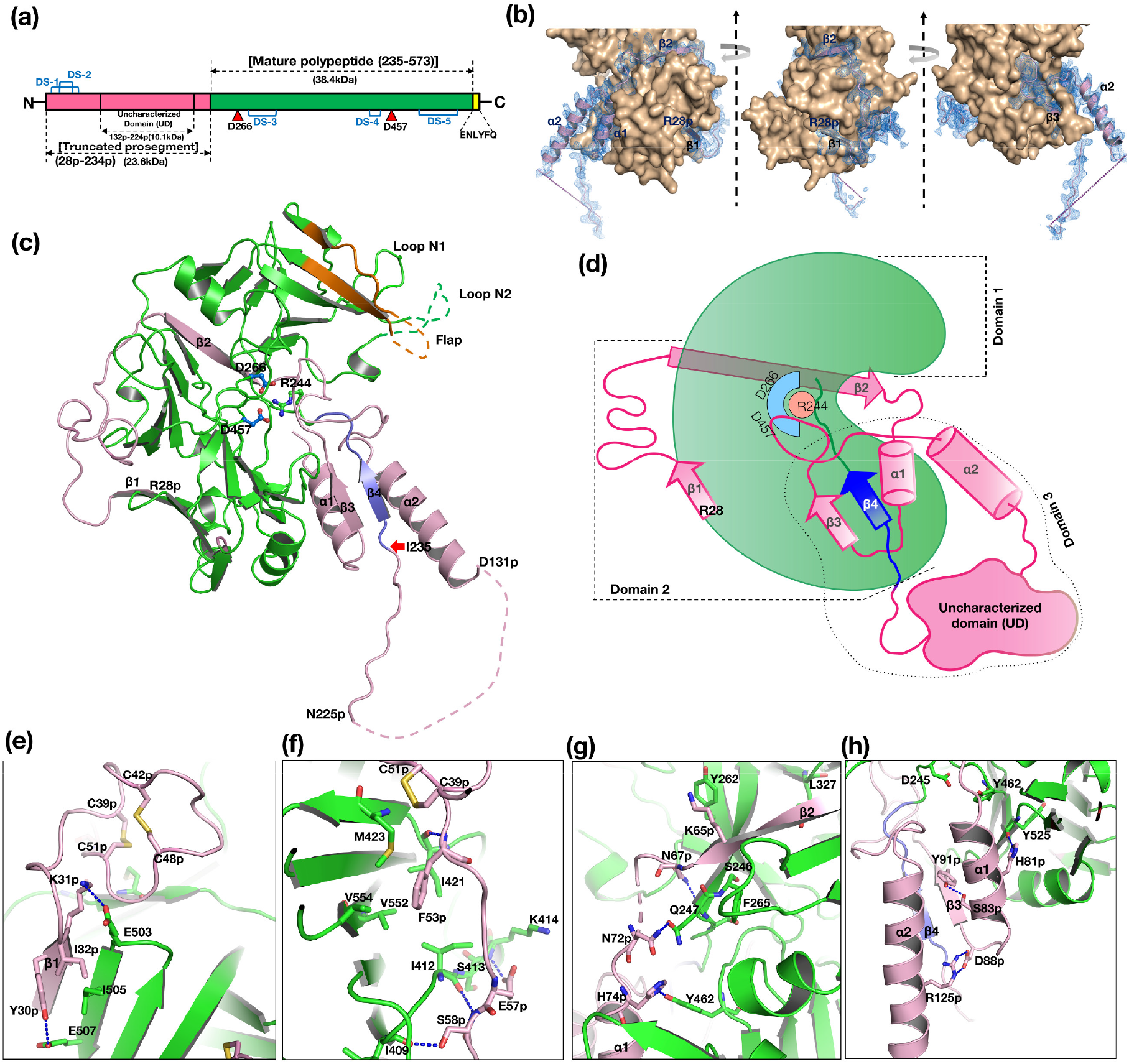
Folding of *Pf*pro-PMX prosegment and its stabilizing interactions with the mature part of the protease. (**a**) Schematic diagram of full-length *Pf*pro-PMX (construct 1) containing truncated prosegment (R28p-N234p; ‘p’ indicates the residues from prosegment) with uncharacterized domain (UD, E132p-K224p) in pink, mature part in green (I235-N573) and TEV protease site at the C-terminus in yellow. The disulfide bonds are shown with blue bridges and catalytic aspartates with red triangles. (**b**) *2F*_*o*_*-F*_*c*_ electron density (blue mesh, 1σ contour level) of *Pf*pro-PMX prosegment (light pink cartoon) wrapped around mature protease (brown surface) shown by views rotated by 90°. The N-terminal residue R28p and secondary structure elements of the prosegment have also been marked and no density observed for UD (dotted line). (**c**) Overall structural fold of *Pf*pro-PMX in cartoon representation, with prosegment (light pink), the mature region (green) and flap (orange), the first β-strand of mature segment in blue and the missing UD dashed lines. The catalytic aspartates (blue carbon) and inhibitory arginine (R244, green carbon) are shown, along with the red arrow indicating the putative first residue of the mature enzyme. Two loops (Loop N1 and N2) in the mature region are marked. (**d**) Schematic diagram of *Pf*pro-PMX showing the domains and the secondary structural elements of prosegment (pink) around the mature protease (green). The first β-strand of the mature segment in blue followed by a loop region in green color. (**e-h**) The interactions of the prosegment (pink) with the mature region (green) of *Pf*pro-PMX.

The overall fold of the prosegment of *Pf*pro-PMX (**Fig. 1c**) is unique and such an arrangement of the prosegment has not been observed before for any other zymogens of pepsin-like aspartic proteases. The prosegment wraps around the elongated bean-shaped mature enzyme (**Figs. 1c and 1d**). The structure of *Pf*pro-PMX can be divided into three segments: domain 1 (F248 to D391), domain 2 (R28p-K68p and P392-K572), and domain 3 (P71p to Q247) (**Fig. 1d**). Domain 3 also consists of an “uncharacterized domain” (UD) which contains ∼90 residues (E132p-K224p) and is not visible in the crystal structure. The prosegment is folded into the first β-strand (β1, R28p I32p), followed by an extended loop, the second β-strand (β2, S59p-L66p), an α-helix (α1, H74p-K85p), the third β-strand (β3, D88p-N94p), a second loop with a twisted conformation, the second long α-helix (α2, M114p-D131p), and finally an extended loop that connects to the mature part of the protease. The prosegment is stabilized by the mature domain of *Pf*pro-PMX through several polar and hydrophobic interactions (**Figs. 1e-1h**). Additionally, the curved conformation of the first loop region between β1 and β2 of the prosegment is stabilized by two disulfide linkages formed between C39p and C51p, as well as C42p and C48p (**Extended Data Fig. 1b**). Part of the prosegment (R95p-G104p) is folded in a unique twisted conformation (twisted loop), which then blocks the S1’ and S2’ pockets of the active site (**Fig. 2a**). The C-terminal part of the twisted loop surrounds the initial segment (P239-K241) of the mature polypeptide. A salt bridge between the side chains of R95p and E433 stabilizes the initial part of this loop. The side chains of I99p and L103p are packed in the hydrophobic pocket formed by the flap region of the mature segment (**Fig. 2b**).

**Figure 2.**
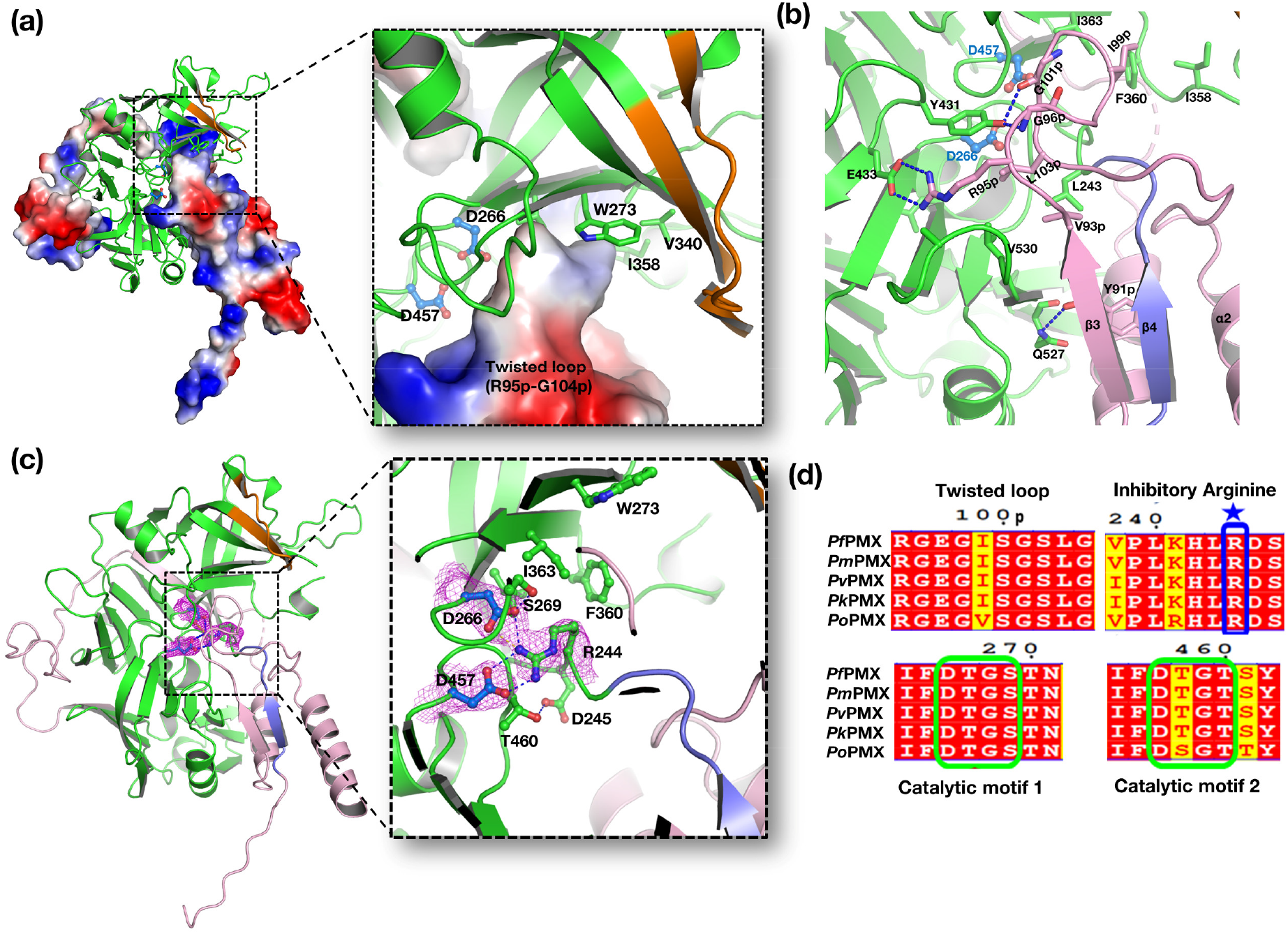
Blocking of *Pf*pro-PMX active site by the twisted loop of the prosegment and inhibitory arginine (R244). (**a**) The prosegment is shown as an electrostatic surface (blue: positive, red: negative and white: neutral) and the mature part of the enzyme in green and flap is in orange cartoon representation. The inset shows the zoomed in-view of the twisted loop blocking the active site. (**b**) The twisted loop region blocks the active site residues (D266 and D457, blue ball-and-stick). The β3 and following twisted loop region are held by multiple interactions. (**c**) The structure of *Pf*pro-PMX highlighting blockage of the catalytic aspartates (D266 and D457) by R244 mediated through salt bridge interactions. Inset shows *2F*_*o*_*-F*_*c*_ electron density map contoured at 1σ level around R244, D266 and D457. The other neighbouring residues of catalytic aspartates are also shown. (**d**) Conservation of the residues of the twisted loop, inhibitory R244 and the catalytic motifs in PMX zymogens from other human malaria causing *Plasmodium* species [*P. falciparum* (Pf), *P. vivax* (Pv), *P. knowlesi* (Pk), *P. ovalae* (Po)].

The first five residues (I235-P239) of the mature polypeptide form a β-strand (β4) parallel to the β3 of the prosegment. Further downstream the mature peptide is extended into the active site that is stabilized by several interactions with the main chain and side chain atoms. One notable interaction in the initial part of the mature region is formation of a salt bridge by R244 with the carboxylate groups of the catalytic aspartates (D266 and D457) (**Fig. 2c**). Sequence comparison also reveals that R244 is highly conserved in PMXs from other malaria-causing parasites (**Fig. 2d**). The flap of the zymogen is in an open conformation due to folding of the twisted loop of the prosegment that occupies the active site region. Two other loops, N1 and N2, also assume an open conformation, likely due to the folding of the polypeptide that connects the second β-strand and the first α-helix of the prosegment (**Fig 1c**). Apart from the two disulfide linkages in the prosegment, *Pf*pro-PMX structure has three more disulfide bridges in the mature domain (**Extended Data Fig. 1b**). Two disulfide bonds (C279-C284 and C482-C521) are similar to those observed in structures of other plasmepsins and pepsin-like aspartic proteases. The disulfide bond formed by two adjacent cysteine residues (C447-C448), present in the *Pf*pro-PMX structure, has not been previously reported for any other pepsin-like aspartic proteases. Sequence alignment shows that five disulfide bonds observed in the *Pf*pro-PMX structure should be conserved in other PMXs from parasites that cause malaria in humans (**Extended Data Fig. 2a**).

Analysis of the arrangement of the protein molecules in the crystals reveals that *Pf*pro-PMX forms an “S” shaped dimer, also observed in zymogens of other vacuolar PMs^22^. The loop region of the prosegment consisting of residues N225p-K232p is swapped between the two adjacent monomers of the dimer (**Extended Data Figs. 2b and 2c**). Five residues (T227p-T231p) of one monomer form a β-strand that results in the formation of an anti-parallel β-sheets by pairing with the first β strand of the mature polypeptide of another protomer. It is not clear whether such a dimer would form in presence of the UD, which in the current structure is not visible. The UD from different PMXs exhibit variable lengths and very low sequence similarity. However, the sequences without UD are highly conserved, indicating that the PMX zymogens from different malaria-causing parasites might have similar overall structures, with the twisted loop from the prosegment and arginine (R244) from the mature peptide blocking the active site (**Extended Data Fig. 2a**).

### Structural comparison of *Pf*pro-PMX with zymogens of other pepsin-like aspartic proteases

*Pf*pro-PMX has been compared with the representative zymogen structures of other pepsin-like aspartic proteases from parasites, plants, and animals. Previously reported crystal structures of the truncated zymogens of *P. falciparum* vacuolar plasmepsins (*Pf*PMII, *Pf*HAP, and *Pf*PMIV) reveal that their prosegments have an initial β-strand, followed by two α-helices connected by a turn and a coiled extension with the characteristic “Tyr-Asp” loop that connects to the mature segment of the enzyme^22^. The prosegment of the plant and gastric pepsin-like aspartic protease zymogen comprises the first β-strand, followed by two α-helices, a short α-helix, and a turn connecting to the mature domain of the enzyme.

Superposition of the *Pf*pro-PMX structure with the structures of pepsinogen (3PSG, representing plant and gastric pepsin-like aspartic protease zymogens) and *Pf*PMII zymogen (1PFZ, representing vacuolar PM zymogens) shows that the folding of its prosegment is remarkably different as compared to the latter structures (**Fig. 3a, Supplementary movie 2**). *Pf*pro-PMX has a much longer prosegment than the other pepsin-like aspartic protease zymogens (**Extended Data Fig. 1a**). The β2 of *Pf*pro-PMX occupies the same position as the first β-strand of pepsinogen and *Pf*PMII zymogen structures (**Fig. 3b**) and the remaining secondary structure elements have different orientations. The α1 of *Pf*pro-PMX points downwards, occupying the same space as the tip of a loop in *Pf*PMII zymogen (115p-126p) and pepsinogen; hence similar loop is absent in the *Pf*pro-PMX structure. This loop of vacuolar PM zymogens plays an important role in their activation mechanism^22^; the absence of such structural element in *Pf*pro-PMX may imply a different mechanism of maturation. The next secondary structural elements in pepsinogen and *Pf*PMII zymogen are α-helices pointing in two different directions; in *Pf*pro-PMX it is a β-strand. The last part of the prosegment in pepsinogen forms a short helical turn that provides a lysine residue blocking the active site aspartates; an equivalent position is partly occupied by the twisted loop region of *Pf*pro-PMX. In vacuolar PM zymogens the second α-helix is involved in formation of the inter-dimer contacts and the “Tyr-Asp” loop keeps the prosegment in a strained conformation^22^. Inactivation of vacuolar PM zymogens is achieved by the prosegment that acts as a harness separating the two domains of the enzyme, resulting in distancing of the catalytic aspartates. High sequence conservation of the twisted loop region (R95p-G104p) among PMX zymogens from other malaria-causing parasites (**Fig. 2d**) implies homology in their structural features. The sequence and phylogenetic analysis (**Supplementary Figs. 2a and 2b**) of the UD reveals that this region might be of plant origin and its structural superposition shows that this region may occupy a similar space to that observed for the plant specific insert in phytepsin zymogen structure (1QDM). However, only determination of the crystal structure of *Pf*pro-PMX with this domain visible would confirm the actual fold and 3D arrangement of this insertion region. The pro-mature cleavage site (between N234p and I235) of *Pf*pro-PMX is distal from the active site as seen in vacuolar plasmepsin zymogens, whereas the cleavage site of pro-mature region is adjacent to the catalytic aspartates in the zymogens of plant and gastric pepsin-like aspartic proteases.

**Figure 3.**
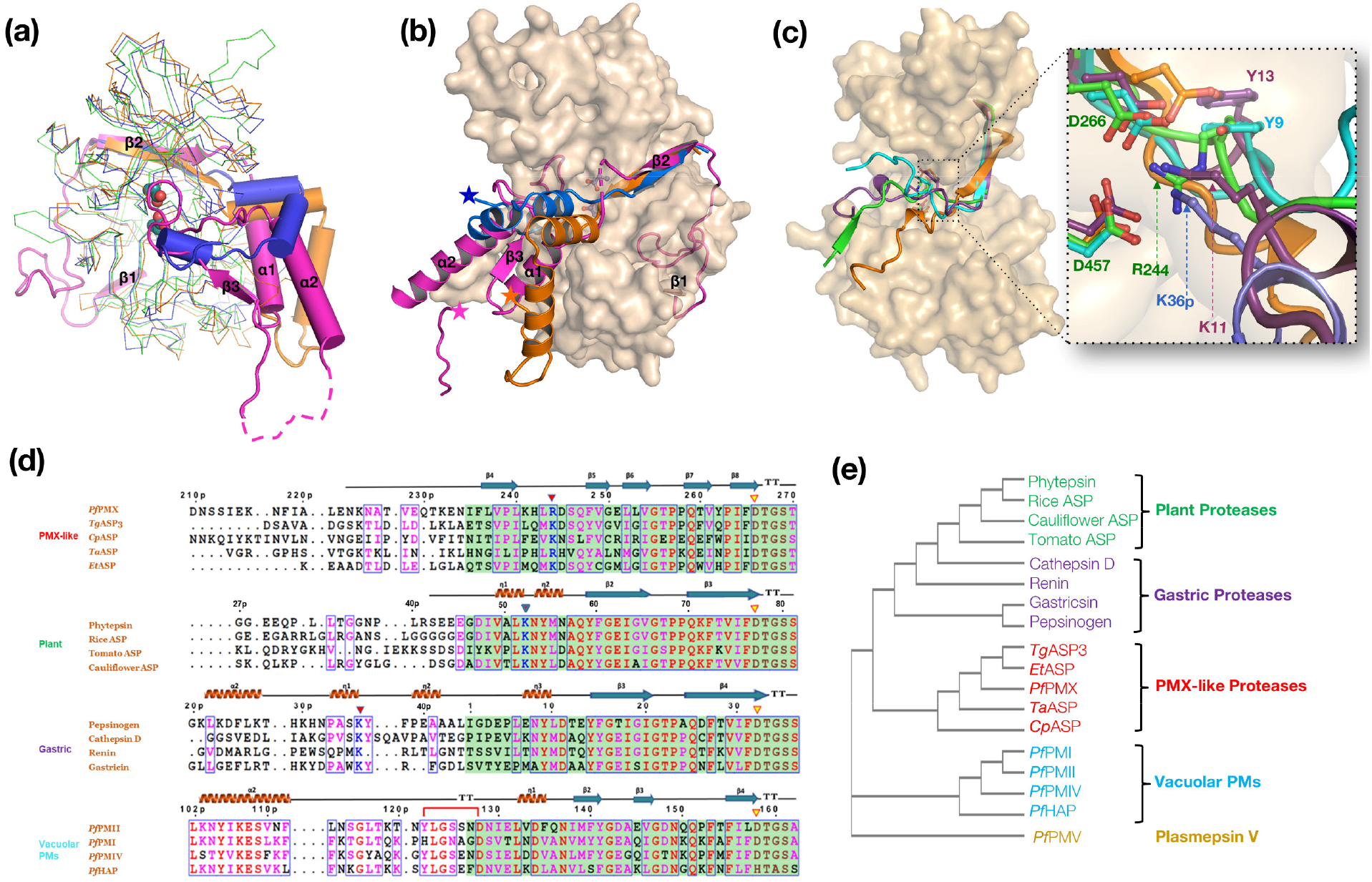
Comparison of *Pf*pro-PMX with vacuolar PMs, plant, and gastric pepsin-like aspartic protease zymogens. (**a**) Superposition of *Pf*pro-PMX (magenta prosegment and green mature enzyme) structure with pepsinogen (blue, 3PSG), and *Pf*PMII zymogen (orange, 1PFZ). The mature enzymes are shown in thin ribbon, prosegments in cartoon, the catalytic aspartates by cyan sphere, and the secondary structural elements of *Pf*pro-PMX prosegment are labelled. (**b**) The comparison of the prosegment fold of *Pf*PMX (magenta), *Pf*PMII (orange), and pepsinogen (blue). The mature part of *Pf*pro-PMX is shown as surface and stars represent the end of the prosegment of different zymogens. (**c**) The comparison of the initial segments of the mature peptides of *Pf*pro-PMX (green), *Pf*PMII zymogen (orange), pepsinogen (cyan), and phytepsin zymogen (violet). The residues occupying the active site and catalytic aspartates are shown in sticks. Inset shows the zoomed in-view indicating the blockage of the active site resulting in the inactivation of the enzyme. The prosegment of the pepsinogen is shown in blue. The residues are numbered as they are assigned in their corresponding structures in PDB. (**d**) Multiple sequence alignments showing the last part of the prosegment and initial part of the mature segment of the zymogens of PMX-like proteases, plant and gastric pepsin-like aspartic proteases as well as vacuolar PMs. The mature region is highlighted with a green background. Sequence numbering is given on top of each alignment, ‘p’ refers to the residues from prosegment. The inhibitory lysine/arginine residues (blue) and the catalytic aspartates (brown) are marked with a red and yellow triangle, respectively. The Tyr-Asp loop in vacuolar PMs is shown by bridged red line. Sequences used: *Plasmodium falciparum* (*Pf*PMX, XP_001349441.1), *Toxoplasma gondii* (*Tg*ASP3, XP_002367043.1), *Cryptosporidium parvum* (*Cp*ASP, XP_627444.1), *Theileria annulata* (*Ta*ASP, XP_954092.1), *Eimeria tenella* (*Et*ASP, XP_013233972.1), Phytepsin (KAE8806867.1), Rice ASP (BAA0687.5), Tomato ASP (NP_001234702.1), Cauliflower ASP (CAA56373.1), porcine pepsinogen (AAA31096.1), Cathepsin D (AAp36305.1), Renin (XP_003823019.2), Gastricin (NP_002621.1), *Pf*PMII (XP_001348250.1), *Pf*PMI (XP_001348249.1), *Pf*PMIV (XP_001348248.1), *Pf*HAP (XP_001348251.1). (**e**) Phylogenetic tree developed from the full-length sequence alignment of different pepsin-like aspartic protease zymogens.

A sequence alignment and superposition of homologous modeled structures of other PMX-like protease zymogens from different apicomplexan parasites reveals that these proteases should also assume similar structural folds, with two characteristic disulfide bridges in the prosegment along with the blocking of the active site by a twisted loop (**Extended Data Figs. 3a and 3b**). The folding of the initial segment of the mature peptides in the zymogen structure of pepsin-like aspartic proteases are quite different (**Fig. 3c**). The first part (I235-K241) of the mature polypeptide of *Pf*pro-PMX structure forms a β-strand parallel to the β3 of the prosegment. The polypeptide that connects the first and second β-strands of the mature enzyme is extended, with R244 forming salt bridge interactions with the two catalytic aspartates (**Fig. 2c**). PMX-like proteases from other parasites also have arginine or lysine at the same position, forming salt bridge interactions with the catalytic aspartates (**Extended Data Figs. 3a and 3b**). Blocking of the catalytic aspartates by R244 from the mature polypeptide is unique to *Pf*pro-PMX. Although a lysine residue in the phytepsin zymogen (K11 from mature polypeptide, 1QDM) and pepsinogen (K36p from prosegment, 2PSG) is blocking the catalytic aspartates, there is an additional tyrosine residue (Y13 in 1QDM; Y9 in 2PSG) which also forms a hydrogen bond with the catalytic residues (**Figs. 3c and 3d, Supplementary movie 3**), but the latter is absent in PMX-like proteases or in vacuolar PM zymogens. Further analysis of the full-length sequence based structural alignment of different pepsin-like aspartic protease zymogens also places PMX-like proteases close to plant and gastric proteases (**Fig. 3e**).

### Modeling of the structure of mature *Pf*m-PMX and its complex with a substrate

A model of the mature enzyme, *Pf*m-PMX (**Fig. 4a**) was developed. The accuracy and the stability of *Pf*m-PMX structural fold are supported by MD simulation of the structures with and without inhibitors (**Extended Data Figs. 4a-d**). *Pf*m-PMX has pepsin-like aspartic proteases fold with two topologically similar N- and C-terminal domains, but with some striking differences as compared to the enzymes belonging to same family. The N-terminal domain has a β-hairpin-like structure referred to as the flap (I303-T322). One notable substitution in *Pf*m-PMX is F311, located in the tip of the flap. An equivalent position in other pepsin-like aspartic proteases (except HAP) is occupied by a conserved tyrosine residue (Y77 in pepsin) which, upon substrate binding, forms a hydrogen bond with a tryptophan residue (W41 in pepsin) due to flap closure, thus initiating a proton relay to maintain the protonated state of the N-terminal catalytic aspartate^25^. Presence of F311 in the case of *Pf*m-PMX would change the opening and closing dynamics of flap, implying a modified mechanism of catalysis without the assistance of a tyrosine. The mature part of *Pf*m-PMX has three disulfide bonds between residues C279-C284, C447-C448, and C482-C521; whereas vacuolar plasmepsins usually have only two disulfide bonds (C45-C50 and C249-C282 in *Pf*PMII). Due to formation of the disulfide C482-C521 in *Pf*m-PMX, the peptide region from residues H478-K492 which has a short loop and helical conformation is stabilized (**Supplementary Fig. 3a**). An equivalent disulfide bond is not observed at same location in other pepsin-like aspartic proteases.

**Figure 4.**
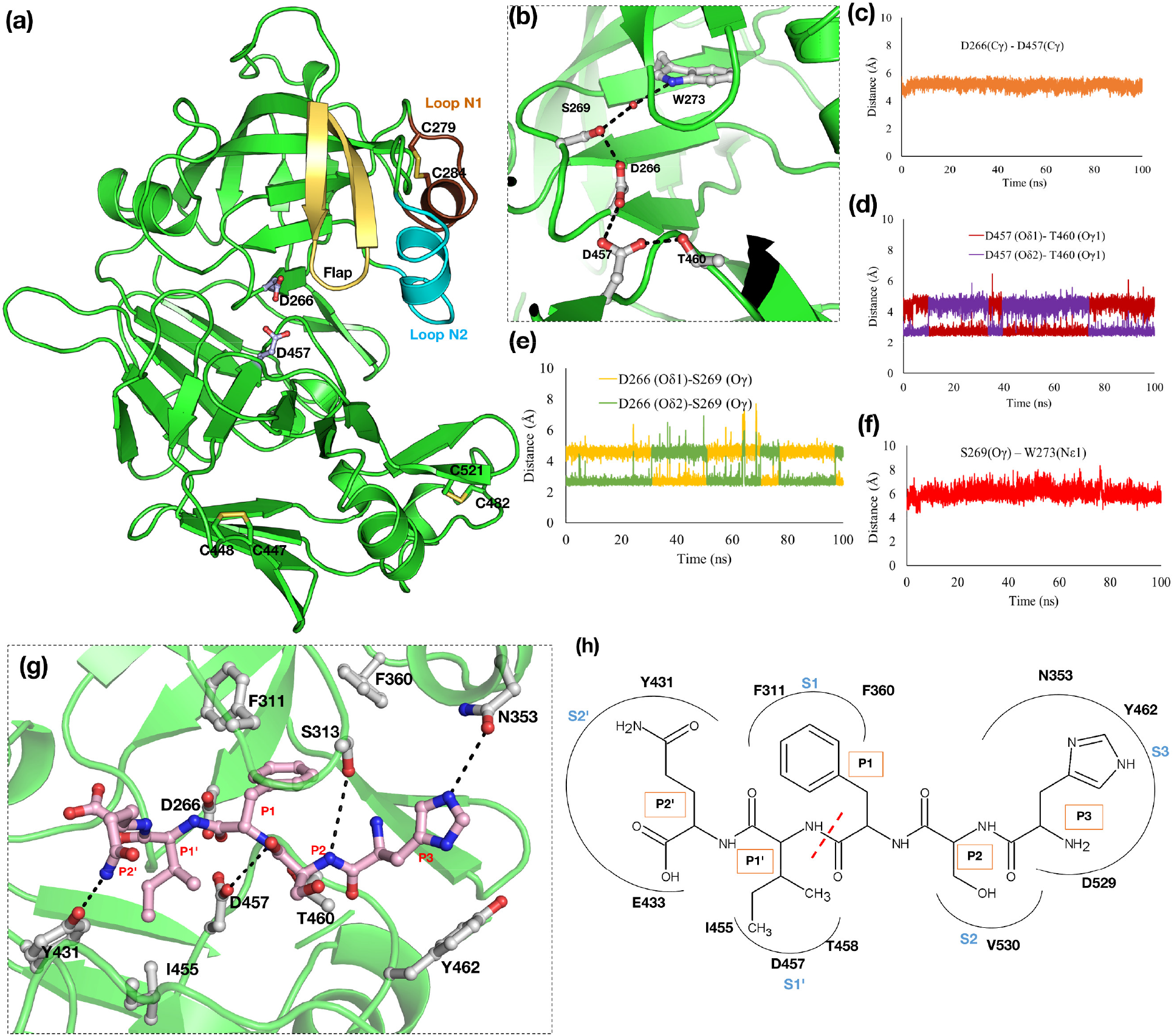
Structural features of Apo- and substrate bound *Pf*m-PMX. **(a)** Cartoon representation of the overall structure of *Pf*m-PMX showing the pepsin-like aspartic protease fold with the catalytic aspartates (grey, ball and stick), flap (golden), disulfide bonds (sticks), Loop N1(brown) and loop N2 (cyan) highlighted. **(b)** The orientation of the key active site residues of *Pf*m-PMX involved in catalysis. The hydrogen bonding network is indicated by the black dashes. The distances between **(c)** D266(Cγ)-D457(Cγ), **(d)** D457(Oδ1-Oδ2)-T460(Oγ1), **(e)** D266(Oδ1-Oδ2)-S269(Oγ), **(f)** S269(Oγ)-W273(Nε1) during the simulation run indicating stability of these interactions in the *Pf*m-PMX-Apo structure. **(g)** Binding mode of peptide (Rh2N, pink) in the active site pocket of *Pf*m-PMX. The residues in substrate binding pocket are in grey and interactions are shown with black dashes. **(h)** Schematic diagram of the substrate binding pocket subsites (S) with corresponding residues of *Pf*m-PMX and its peptide substrate (P) shown with covalent structure. The red dotted line shows the peptide bond hydrolyzed by *Pf*m-PMX.

Analysis of the MD simulation run of the *Pf*m-PMX-Apo provides insights into the active site architecture (**Fig. 4b**) of the enzyme. The Cγ atoms of the carboxylate groups of D266 and D457 are at a distance of around 6.0 Å (**Fig. 4c**), indicating the proximity of these two residues that are needed for catalytic competence. However, carboxylate groups of these two catalytic residues seem to rotate during simulation. The carboxylate group of D457 forms a hydrogen bond with the Oγ1 atom of the T460 side chain (**Fig. 4d**). Also, the Oγ atom of the side chain of S269 is in a constant hydrogen bonding distance from the carboxylate group of the catalytic D266 (**Fig. 4e**) and around 6.0 Å away from the Nε1 atom of W273 (**Fig. 4f**). All these distances are similar to the corresponding distances reported for the crystal structures of other pepsin-like aspartic proteases. The presence of a water molecule near the carboxylate group of D457 during the MD simulation also confirms its ability to act as a base during catalysis, as in other pepsin-like aspartic proteases. The side chains of W273 and S269 form a water-mediated interaction which is also extended to D266 somewhat similar to other pepsin-like aspartic proteases. The structure of *Pf*m-PMX indicates that a proton relay from W273 can occur and aid in catalysis even without the help of the flap tyrosine residue.

To decipher the molecular basis of substrate recognition by *Pf*m-PMX, the peptide Rh2N with the cleavage site (^P3^HSF/IQ^P2’^)^19^ sequence was docked into the protein active site and the complex was simulated. The results indicate that the substrate is oriented in a conformation with the phenylalanine at P1 occupying the S1 hydrophobic pocket formed by the flap as well as by loops N1 and N2 (**Figs. 4g and 4h, Supplementary movie 4**). The amide backbone atom of phenylalanine (P1) is hydrogen bonded to D457 and its carbonyl group is pointing towards D266. The side chain of the serine (P2) points away from the flap and its amide backbone interacts with the side chain of the flap S313. Histidine (P3), the N-terminal residue of the substrate, occupies the S3 pocket of the protein formed by residues N353, Y462, and D529. The Nε2 atom of the P3 histidine interacts with the side chain of N353 of loop N2. The side chain of glutamine (P2’) at the C-terminal end makes an interaction with Y431 (**Figs. 4g and 4h, Supplementary movie 4**). Most of these residues interacting with substrates are conserved among the PMX-like aspartic proteases implying similar mode of substrate binding in these enzymes.

### Removal of the prosegment of *Pf*pro-PMX to form mature *Pf*m-PMX

The full-length polypeptide (∼65 kDa) of *Pf*PMX zymogen is longer compared to other vacuolar PM zymogens (**Extended Data Fig. 1a**), and it also has a UD (10.1 kDa) inserted into the prosegment (**Fig. 1a**). A construct (construct 1, R28p-N573 with prosegment R28p-N234p) of *Pf*pro-PMX, expressed a fusion protein (**Fig. 5a**) with MW ∼62 kDa in Expi 293 GnTI-cells and analysis of the eluted protein fractions shows four distinct bands corresponding to the molecular weights of ∼62 kDa (full length), ∼50.5 kDa, ∼40 kDa and ∼12.1 kDa (**Fig. 5b**). N-terminal sequencing of the ∼40 kDa band indicated that the putative mature domain begins at L221p and the ∼12.1 kDa fragment starts at R28p. Similar pattern of cleavage was also observed when another construct (Construct 2, N190p-N573 with prosegment N190p-N234p) expressed a fusion protein with MW ∼61.2 kDa in *E. coli* (**Supplementary Data 1**). These results suggest that the maturation of *Pf*PMX zymogen occurs via cleavage of the prosegment at multiple cleavage sites (CS-1 and CS-2).

**Figure 5.**
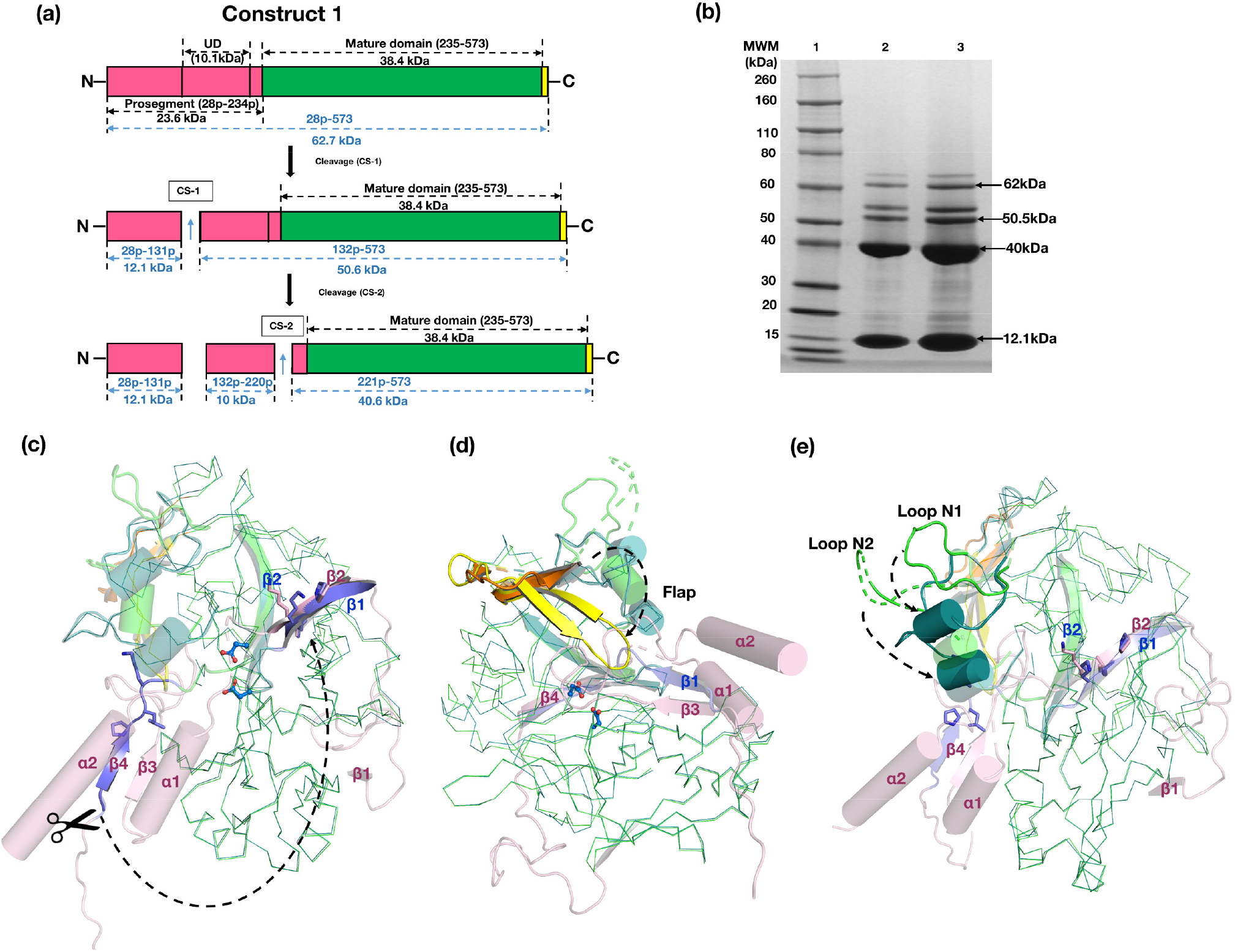
Maturation of *Pf*pro-PMX and associated structural change during the activation process. **(a)** Schematic representation of construct 1 of *Pf*pro-PMX, showing maturation pattern at cleavage site-1 (CS-1) and cleavage site (CS-2) **(b)** SDS-PAGE analysis of the eluted fractions of *Pf*pro-PMX (construct 1) after size exclusion chromatography, lane1: molecular weight marker (MWM), lane2 and 3: eluted fractions. **(c)** Structural changes in *Pf*pro-PMX during maturation process. The secondary structural elements of the prosegment of *Pf*pro-PMX (pink) and first β-strand of the mature (β1, violet) *Pf*m-PMX are shown as a cartoon. β4 (violet) of the zymogen undergoes structural rearrangement and becomes β1 of the mature enzyme; the latter (β1) will occupy the position of β2 of zymogen upon maturation (dotted arrow). Catalytic aspartates are shown as blue colored ball and stick and the possible pro-mature cleavage site is indicated by a scissor. (**d**) Conformational changes (dotted arrow) in flap of *Pf*pro-PMX (orange) and *Pf*m-PMX (yellow). **(e)** Conformational change of Loops N1 and N2 during the maturation of *Pf*pro-PMX (green) and *Pf*m-PMX (teal). For the purpose of visualization, the loop N2 in the *Pf*pro-PMX has been shown as green dotted lines.

Comparison of the crystal structure of *Pf*pro-PMX with the model of *Pf*m-PMX reveals that the maturation process of *Pf*PMX zymogen would occur with significant structural changes in the protein (**Supplementary movie 5**). Upon cleavage of the prosegment, the β4 (I235-K241) of the zymogen folds upward to form the first β-strand (β1) of the mature enzyme. Notably, a tripeptide stretch of β1 of the mature enzyme with the sequence P239-L240-K241 is identical to the sequence P63p-L64p-K65p, which is located in the β2 of the prosegment. This indicates a possibility that the β1-strand (P239-L240-K241) of the mature enzyme will occupy the same position as that of β2-strand (P63p-L64p-K65p) of the zymogen (**Fig 5c**). After the disruption of the salt bridge with the catalytic aspartates, R244 forms a cation-π interaction with the side chain of Y357 from the loop N2, as we observed in our experiment (**Supplementary Fig. 3b**). R244 and Y357 are also conserved, indicating the presence of such interactions in PMXs from other human malarial parasites (**Extended Data Fig. 2a**). Removal of the prosegment creates free space, and, as a result, the flap adopts a closed conformation (**Fig. 5d**) with loops N1 and N2 moving downwards (**Fig. 5e**). Such conformational changes involving these secondary structure elements are also observed in other pepsin-like aspartic proteases. Implications of these regions for inhibitor binding have been discussed in the subsequent sections.

### Mode of binding of 49c in the *Pf*m-PMX active site

The compound 49c (**Fig. 6a**), containing a hydroxylethylamine scaffold, was designed as a peptidomimetic inhibitor of plasmepsins^26^ and is reported to inhibit *Pf*PMX^12,18^. The effect of 49c on PfPMX stability at different pH conditions was monitored using nanoDSF (**Fig. 6b**). Difference in melting temperature (ΔT_m_°C) of Apo- and 49c bound *Pf*PMX suggests that the binding of 49c increases thermal stability of the enzyme. Sequential dip in the difference in melting temperature (ΔT_m_°C) is observed with an increase in pH, with the highest ΔT_m_°C observed at pH 5.5, suggesting that binding of 49c is pH dependent.

**Figure 6.**
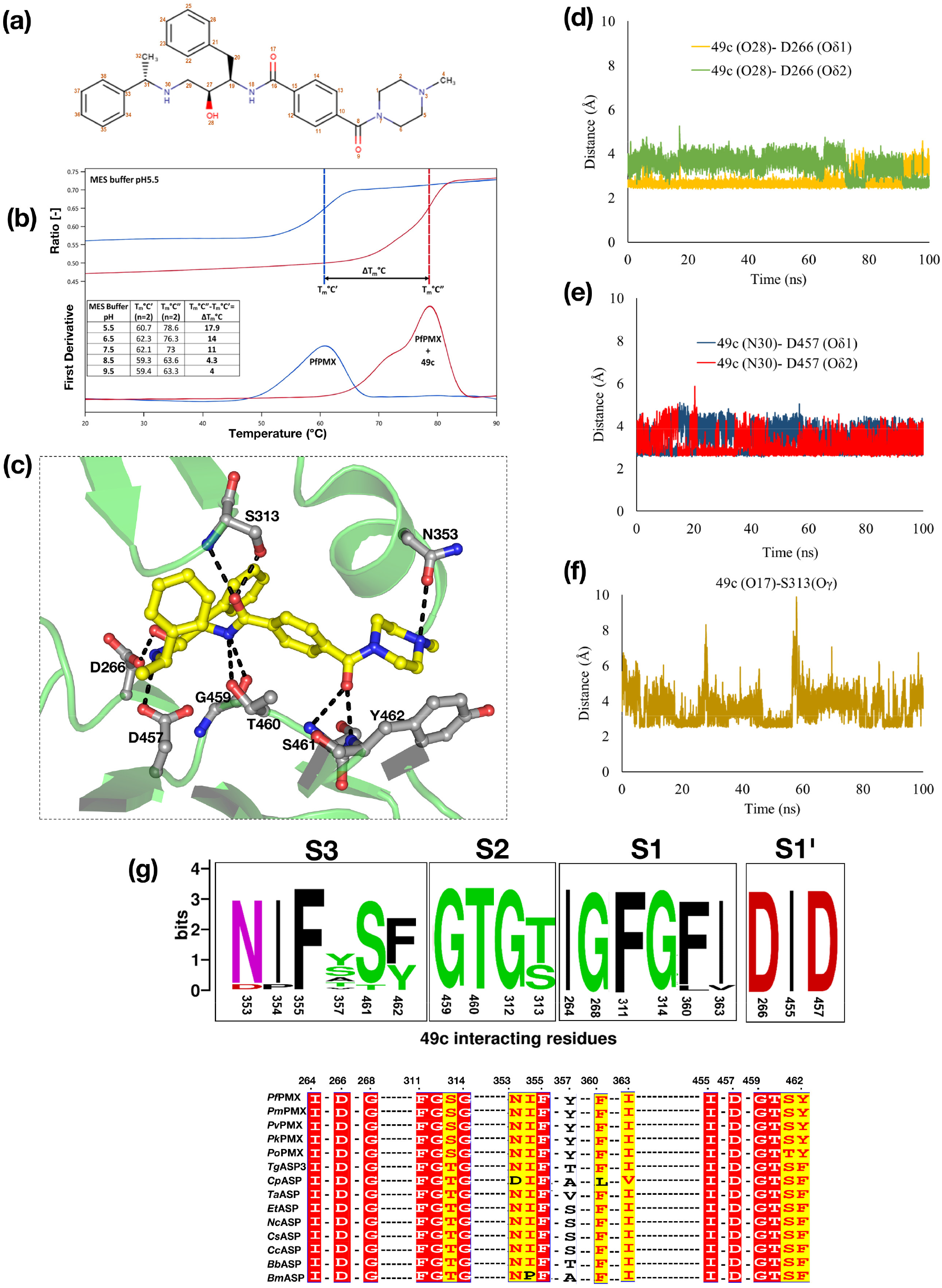
Molecular basis of inhibition of *Pf*PMX by 49c. **(a)** Covalent structure of 49c. **(b)** The change in melting temperature ΔTm°C translates the effect of compound 49c on PfPMX thermal stability. The binding suggested by this effect is pH dependent and it is much stronger in acidic conditions. **(c)** The interactions stabilizing the binding of 49c (yellow carbon) in the active site pocket of *Pf*m-PMX. The hydrogen bonding network is indicated by the black dashes and residues are shown in grey. **(d)** The distances between the central hydroxyl group (atom number O28) of 49c with D266 carboxylate group oxygen atoms. **(e)** The distances between the nitrogen atom (N30) of the main chain NH group of 49c with D457 carboxylate group oxygen atoms. **(f)** The distance between the carbonyl oxygen (O17) atom of 49c with S313 side chain hydroxyl group oxygen (Oγ). **(g)** The conservation and similarities among the residues from PMX-like proteases involved in binding of 49c. The top panel shows the WebLogo (http://weblogo.berkeley.edu) generated using alignment of PMX-like proteases sequences (bottom panel). The letter size in the WebLogo is proportional to residue conservation. The proteases from parasites used in the alignment are: *Plasmodium falciparum* (*Pf*PMX, XP_001349441.1), *Toxoplasma gondii* (*Tg*ASP3, XP_002367043.1), *Cryptosporidium parvum* (*Cp*ASP, XP_627444.1), *Theileria annulata* (*Ta*ASP, XP_954092.1), *Eimeria tenella* (*Et*ASP, XP_013233972.1), *Neospora caninum* (*Nc*ASP, XP_003885934.1), *Cystoisospora suis* (*Cs*ASP, PHJ19229.1), *Cyclospora cayetanensis* (*Cc*ASP, OEH78578.1), *Besnoitia besnoiti* (*Bb*ASP, XP_029222308.1), *Babesia microti* (*Bm*ASP, XP_021337801.1).

To understand the molecular basis of the binding of 49c to *Pf*m-PMX, the compound was computationally docked into the enzyme active site and the inhibitor bound complex was simulated. Here we present, for the first time, a detailed picture of the interactions of 49c with *Pf*m-PMX (**Fig. 6c**). Several polar and hydrophobic interactions stabilize the binding of 49c in the active site of *Pf*m-PMX (**Supplementary movie 6**). The inhibitor binds in a conformation such that its phenylalanine group occupies the S1 pocket, making π-π stacking interactions with F311, similar to those observed for the substrate (**Extended Data Fig. 5a**). The central hydroxyl group of the inhibitor points towards D266. As the phenylalanine group of 49c mimics the P1 group of a substrate, inhibitors with identical or similar groups at this position could possibly have a binding mode similar to 49c or the substrate. The 49c interacts with the catalytic aspartates D266 and D457, as well as with F311 and S313 of the flap. The carboxylate group of D266 is in constant hydrogen bonding with the hydroxyl group (atom number 28) of phenylalanine moiety (**Fig. 6d**) and the carboxylate group of D457 side chain is interacting with the N30 atom of the phenylethylamine moiety of 49c (**Fig. 6e**). Also, the carbonyl group of benzamide (O17) interacts with S313 from the flap during most of the time of simulation, holding the flap in a closed state (**Fig. 6f**). Hydrogen bonding of D266 and D457 carboxylate groups with the hydroxyl groups of the nearby residues S269 **(Extended Data Fig. 5b)** and T460 (**Extended Data Fig. 5c**) respectively, hints at the correct geometry of the catalytic residues in the inhibitor-bound state. Based on sequence analysis, it appears that the residues which are involved in binding of 49c in the active site of *Pf*m-PMX are also conserved or similar (**Fig. 6g**) among PMX-like proteases from various apicomplexan parasites. Therefore, it is likely that 49c would inhibit PMX-like proteases and arrest invasion and egress processes in apicomplexan parasites. Such molecular insights unravelling the interactions of 49c with *Pf*m-PMX would allow development of potent inhibitors for PMX-like proteases.

Although our data suggest that *Pf*m-PMX active site can accommodate inhibitors which are transition state mimics with a central hydroxyl group, notably this enzyme is not inhibited by pepstatin A, a potent inhibitor of a majority of pepsin-like aspartic proteases^12^. Structural superposition of *Pf*m-PMX and pepstatin A-bound *Pf*PMII (1M43) shows that the ligand binding pocket in both proteases are quite dissimilar (**Supplementary Fig. 4**). The presence of Y462 in *Pf*m-PMX, compared to A219 in *Pf*PMII, narrows the binding pocket, leading to possible steric hindrance and blocks the binding of pepstatin A. Screening of pepsin-like aspartic protease inhibitors against *Pf*m-PMX model suggests that the enzyme has higher affinity towards the compounds which inhibit β-secretase (BACE-I) and HIV-1 proteases (**Supplementary data 2**).

## Discussion

Plasmepsin X (PMX) has gained a lot of recent attention as one of the high priority antimalarial drug targets, due to its involvement in the invasion and egress processes of the malarial parasite. PMX-like enzymes are of great interest for antiparasitic drug development as these pepsin-like aspartic proteases are also essential for the lifecycle of many apicomplexan parasites. We have expressed *P. falciparum* PMX zymogen (*Pf*pro-PMX) in HEK cells and produced stable and soluble protein for structural and biochemical studies. Auto-activation of *Pf*pro-PMX is due to multiple cleavage of the prosegment leading to formation of the active *Pf*m-PMX, similar to the sequential cleavage of the prosegment observed for vacuolar PM zymogens^22^. Notably, the purified enzyme shows high thermal stability at acidic pH in the presence of inhibitor 49c, suggesting protonated active site residues facilitating stabilization of the enzyme-inhibitor complex, as observed for other pepsin-like aspartic proteases.

The *Pf*pro-PMX is much longer compared to its counterparts in other vacuolar plasmepsins^27^, as well as in gastric and plant pepsin-like aspartic protease zymogens and exhibit very low sequence similarity with the enzymes belonging to the same family^20,28^. *Pf*pro-PMX has a unique UD in the prosegment. The *Pf*pro-PMX does not have any membrane-anchoring region, in contrast to such regions present in zymogens of vacuolar PMs (PMI-PMIV) and of non-vacuolar PMV^29^. Although the mature *Pf*m-PMX is structurally similar to other pepsin-like aspartic proteases, the folding of the prosegment of *Pf*pro-PMX is unique. Inactivation of *Pf*pro-PMX is not only facilitated by the obstruction of the catalytic aspartates by R244, somewhat similar to the one observed for gastric and plant pepsin-like aspartic protease zymogens which utilize a lysine residue, but also by the projection of the twisted loop region of the prosegment into the active site. Such a mechanism of obstruction of the active site by a prosegment loop region has not been reported so far for any other structurally characterized pepsin-like aspartic protease zymogens. Surprisingly, the inactivation mechanism of *Pf*pro-PMX is vastly different from the mechanism employed by vacuolar PM zymogens.

Inactivation in vacuolar PM zymogens is achieved by spatial separation of the two domains of the mature protein by the prosegment, resulting in the separation of the catalytic aspartates^22,30^, thus rendering the enzyme inactive. Recently, the “S”-shaped dimeric structures of vacuolar PM zymogens have been reported to be associated with their inactive state, while dissociation of dimer to monomer under acidic conditions has been shown to be the first step towards the trans-activation for forming mature enzymes^22^. Analysis of the crystal structure of *Pf*pro-PMX shows the presence of a domain-swapped dimeric structure; however, its implication in inactivation of this enzyme needs further investigations. We suspect that the acidic conditions would cause protonation of the residues of the prosegment of *Pf*pro-PMX resulting in significant structural changes, thus initiating the process of activation. Since, the plausible cleavage sites (CS-1 and CS-2) for activation of *Pf*pro-PMX are far away from the active site, similar to vacuolar PM zymogens, it is likely that the former might also adopt a similar trans-activation pathway as has been recently proposed for the latter^22^. However, the *in-vivo* activation process of *Pf*pro-PMX is ambiguous^24^, hence further studies are needed to provide more insights into the activation mechanism of this enzyme.

After cleavage of the prosegment, the N-terminal polypeptide region of the mature enzyme undergoes significant structural changes that lead to formation of the first β-strand of the active *Pf*m-PMX. Such conformational changes are also observed during the activation process of other pepsin-like aspartic proteases^22,23^. Other notable structural changes of *Pf*pro-PMX include closure of the flap and downward movement of loops N1 and N2. A major substitution in the substrate-binding pocket of *Pf*m-PMX involves F311 from the flap, which replaces the position of the highly conserved Y77 (pepsin numbering) of the pepsin-like proteases. That residue is believed to facilitate the catalytic mechanism by increasing the acidity of the N-terminal catalytic aspartate (D32 in pepsin)^25^. However, in *Pf*m-PMX the proton relay is still possible from W273 to D266 through a water molecule and S269, without the involvement of any residue from flap. Such mechanism is not common among pepsin-like aspartic proteases, although the existence of such a mechanism was postulated for HAP^31^. Notably, strict conservation of F311 in *Pf*m-PMX and in related proteases points towards a possibility of a modified catalytic mechanism. Rotation of the carboxylate groups of the catalytic aspartates (D266 and D457) is observed in a simulated structure of *Pf*m-PMX-Apo and similar multiple conformational states have also been reported for *Pf*PMV^29^ and the crystal structures of *Pf*PMII^32^. Comparison of the P3-P2’ cleavage sites in a majority of the *Pf*m-PMX substrates suggests that the P1 and P1’ are occupied by non-polar residues, whereas P2’ position is mostly occupied by a negatively charged residue. Modifying the P2-P2’ positions to Ala leads to significant loss of cleavage efficiency of *Pf*PMX^19^. The residue at the P1 position is placed in the S1 hydrophobic pocket of the active site of *Pf*m-PMX, similarly to other pepsin-like aspartic proteases^28,29^. A previous report on PMX-like proteases showed the binding of a substrate in the active site of *Pf*PMIX, *Pf*PMX, and *Tg*ASP3 in an opposite orientation as compared to the inhibitor 49c, hence resulted in assignment of the S1pocket as S1’ ^18^. P1 position of the substrate (HSF/IQ) in the *Pf*m-PMX active site in our study follows the same orientation as that depicted earlier for 49c in the *Pf*m-PMX and *Tg*ASP3 active sites. Also, P1’ Ile of our substrate occupies the same position as L103p of the twisted loop in the zymogen, supporting further the mode of substrate binding presented here. Additionally, the two catalytic aspartate carboxylate groups of *Pf*m-PMX are at a favorable distance from the scissile peptide bond (between F and I) to facilitate catalysis. Further detailed structural, biochemical. and quantum mechanical (QM)/molecular mechanics (MM) studies would provide more evidence towards the catalytic mechanism of *Pf*m-PMX.

Despite setting up a variety of crystallization screens, the *Pf*m-PMX-49c complex could not be crystallized. However, the details of the interactions involved in binding of this potent inhibitor to the *Pf*m-PMX active site were obtained from our MD simulation studies. Although the orientation of 49c in the *Pf*m-PMX active site is similar to that reported previously^18^, there are several striking differences observed in our complexed structure and those interactions might be essential for stronger binding of this inhibitor. Docking-based screening highlighted that the inhibitors of BACE-I bind more tightly to *Pf*m-PMX active site. Our structural analysis also provides rationale for the earlier observation that *Pf*m-PMX is not inhibited by pepstatin A, a potent inhibitor of pepsin-like aspartic proteases^12^. Inability of pepstatin A to inhibit *Pf*m-PMX is primarily due to the presence of Y462, which prevents binding of the inhibitor in the S4 pocket of the active site. Similar blocking of the S4 pocket by F370 in *Pf*PMV has been shown to be the primary reason for its lack of binding with pepstatin A^29^.

In conclusion, the experimental structure of *Pf*pro-PMX and computational model of *Pf*m-PMX (both in the apo- and in complex with substrate and inhibitors) provide comprehensive insights into the folding of the prosegment and the architecture of the active site of this enzyme. These findings can be extrapolated to other PMX-like proteases of apicomplexan parasites. The folding of the prosegment, and thus the inactivation mechanism of *Pf*pro-PMX, is novel and, based on sequence and structural similarities, a similar mode of inactivation is anticipated to be present for other PMX-like proteases. As the binding pocket residues are highly conserved among the PMX-like aspartic proteases, the observed mode of binding of 49c and top-hit inhibitors from our computational study in *Pf*m-PMX should aid in screening more drug analogues to identify lead candidates for the development of more potent inhibitors of these classes of enzymes. Therefore, the structural features of the *Pf*m-PMX highlighted in this study should facilitate development of the antiparasitic drugs.

## Materials and Methods

### Expression and purification of recombinant *Pf*pro-PMX (R28p-N573) (Construct 1) in HEK cells

The synthetic gene encoding the *Pf*pro-PMX (R28p-N573) of *Plasmodium falciparum* plasmepsin X (XP_001349441) (**Fig. 5a**) was engineered for expression in a mammalian host (Human Embryonic Kidney cells, HEK cells) by GeneArt (Thermo Fisher, Waltham, MA, USA) and cloned into a variant of pTT5 with the mouse Ig secretion signal peptide on the N terminus and a TEV-cleavable 8x Histidine tag on the C terminus. The recombinant protein was expressed in Expi 293 GnTI- (Thermo Fisher, Waltham, MA, USA) at 37°C, using Expi293 expression media (Thermo Fisher, Waltham, MA, USA). After 5 days of culture, the medium was clarified by centrifugation, filtered, and applied to 2x 5ml HisTrap Ni Excel columns equilibrated with Buffer A (25 mM Tris-HCl, pH 8.0, 200 mM NaCl, 1 mM TCEP, 50 mM Arginine, 0.25% Glycerol). After the protein was loaded, the column was washed with 50 ml of 2% Buffer B (25mM Tris HCl, pH 8.0, 200 mM NaCl, 1 mM TCEP, 500 mM imidazole), followed by 50ml of 4% Buffer B. The protein was eluted with a linear gradient of 4-60% Buffer B. Fractions were analyzed by SDS-PAGE using 4-12% gradient gels in MOPS buffer. Fractions containing the *Pf*pro-PMX were pooled and dialyzed overnight at room temperature into Buffer A containing 10 mg/ml TEV and 170,000 U of EndoH. The extent of tag removal and deglycosylation was assessed by SDS-PAGE. The untagged and deglycosylated protein was applied to 1x 5ml HiTrap Ni Chelating Column equilibrated with Buffer A. The column was washed 50 ml of Buffer A and the protein was eluted with a linear gradient of 0-60% Buffer B. Fractions containing the untagged and deglycosylated *Pf*pro-PMX were observed in the flow thru and fraction 1 of the gradient elution. Fractions were pooled and concentrated to ∼12 mg/ml for size exclusion chromatography (SEC). The concentrated protein was applied to a 1x 120 ml Superdex 200 column in SEC buffer (25 mM Tris-HCl, pH 8.0, 200 mM NaCl, 1 mM TCEP, 1% glycerol). Fractions containing the protein of interest were pooled and concentrated to 11.4 mg/ml for crystallization and a second pool of protein, concentrated to 13.3 mg/ml, was used for assays.

### Study of 49c binding by nanoDSF

A label-free nanoscale Differential Scanning Fluorimetry (nanoDSF) method, was used to assess the effect of compound 49c on *Pf*PMX stability. *Pf*PMX was diluted to 0.5 mg/ml (7.9 µM) in MES buffer in varying pH conditions (25 mM Tris/HCl, 25 mM MES pH 5.5, 6.5, 7.5, 8.5 and 9.5). Thermal stability of *Pf*PMX was characterized at each pH condition with and without 49c at 160 µM. All samples were tested at the final 0.16% concentration of DMSO. Samples were loaded in standard capillaries (PR-C002, NanoTemper Technologies) with a final volume of 10 µl. *Pf*PMX intrinsic fluorescence (aromatic residues fluorescence) was measured over a linear thermal gradient 20°C-95°C at 1°C min^-1^ with an excitation power set to 40%. Fluorescent measurements were made at excitation 280 nm (UV-LED) and at emission wavelengths of 330 and 350 nm using Prometheus NT.48 (NanoTemper Technologies). Data from duplicate measurements of each sample were processed using PR.ThermControl Software (NanoTemper Technologies).

### Crystallization, diffraction data collection and structure solution of *Pf*pro-PMX

Purified *Pf*pro-PMX (SSGCID batch ID PlfaA.17789.b.HE11.PD28363) at 11.4 mg/ml was crystallized via vapor diffusion using a condition derived from JCSG+ screen (RigakuReagents), condition A7 (100mM TrisHCl/NaOH pH 9.4, 18% (w/V) PEG 8000) with 0.4 µl protein + 0.4 µl reservoir as sitting drops in 96-well XJR trays (Rigaku Reagents) at 14°C. Crystals were cryo protected with 20% ethylene glycol as cryoprotectant added to reservoir solution. Diffraction data were collected at the Life Sciences Collaborative Access Team beamline 21-ID-F at the Advanced Photon Source, Argonne National Laboratory using a Rayonix MX-300 detector. The diffraction data were reduced with XDS/XSCALE^33^ to 2.1Å resolution. The intensities were converted to structure factors with the program modules F2MTZ and CAD of CCP4^34^.

The structure was solved via Molecular Replacement with MR-ROSETTA^35^ using PDB entry 3PSG as the search model. The structure was further modeled in iterative cycles of real space refinement in Coot^36,37^, and through reciprocal space refinement in phenix.refine^38^. The structure was subsequently refined for ten cycles using REFMAC5^39^. The resulting electron density map visualized in Coot indicated better features of the missing residues and ligands of the *Pf*pro-PMX model. Subsequently, manual model building was performed by visual inspection and refinement using REFMAC5. The ligands, solvent molecules and ions were progressively added at peaks of electron density higher than 3σ in sigma-A weighted *F*_*o*_*−F*_*c*_ electron density maps while monitoring the decrease of *R*_free_ and improvement of the overall stereochemistry. The polypeptide region with residues K69p-N70p and E132p-K224p in the prosegment and the C-terminal linker region could not be modeled due to the lack of features in the electron density. With the exception of a few residues (H308-S315) in the flap, (E343-D352) in N2 loop and the last residue N573, the complete mature enzyme portion of the *Pf*pro-PMX structure could be modeled and refined. The side chains of the residues with weak or missing electron density have not been added in the structure. The adjacent disulfide bond (447-448) is modelled as *cis* peptide bond conformation^40^. The data collection and refinement statistics are presented in Table 1. Quality assessment tools built into Coot, phenix, and Molprobity^41^ were used to assess the quality of the structure during refinement. The structure was deposited in the PDB with code 7RY7. Diffraction images were made available to the Integrated Resource for Reproducibility in Macromolecular Crystallography (www.proteindiffraction.org)^42^.

### Modeling of mature *Pf*m-PMX

A model of *Pf*m-PMX (I235-K572) was generated by comparing the crystal structure of *Pf*pro-PMX and the model obtained from SwissModel (https://swissmodel.expasy.org/). The quality of the model was assessed by the Ramachandran plot and visual inspection of the active site. Further, the stability of the *Pf*m-PMX model was monitored by molecular dynamics (MD) simulation as discussed in the next section.

### Molecular dynamics (MD) simulations of *Pf*m-PMX as Apo and its complexes

All the molecular dynamics (MD) simulations in this study were performed by GROMACS 2019.1 with ‘Amber99SB’ force field^43^. The force field parameters for the ligands were generated using Antechamber. The ‘*xleap*’ module of AmberTools was used to generate topologies for *Pf*m-PMX Apo as well as complexes taking care of the disulfide bonds. The protein molecules (as apo or complexed) were placed at the center of the box with a distance of 20 Å from the wall surrounded by TIP3P water molecules with periodic boundary condition. The complexes were neutralized by adding the appropriate number of required counter ions. The ParmED (https://parmed.github.io/ParmEd/html/parmed.html) tool was used to convert the topology files obtained from xleap module of AmberTools for the use of those by Gromacs. Energy minimizations were performed for 50,000 steps. The energy minimized systems were further equilibrated using canonical ensemble (NVT) followed by isothermal-isobaric ensemble (NPT). In the NVT equilibration, systems were heated to 300 K using V-rescale, a modified Berendsen thermostat for 1 ns. In NPT, all these heated systems were equilibrated using the Parrinello Rahman barostat for 1 ns to maintain a constant pressure of 1 bar. The unrestrained production MD simulations were performed for 100 ns for all structures. The covalent bonds involving H-atoms were constrained using the ‘LINCS’ algorithm and the long-range electrostatic interactions with particle mesh Ewald (PME) method. The time step for integration was set to 2 fs during the MD simulation.

MD simulation trajectories were further analyzed by using the inbuilt tools in GROMACS 2019.1 and visual molecular dynamics (VMD)^44^. The images were generated using the Discovery studio visualizer (https://www.3ds.com/products-services/biovia/) and PyMol (https://pymol.org/2/). All simulations were performed on the Spacetime High Performance Computing (HPC) resource at IIT Bombay.

### Sequence Alignment

All the sequence alignment has been performed using PROMALS3D multiple sequence and structure alignment server (http://prodata.swmed.edu/promals3d/promals3d.php) and T-COFFEE server (https://www.ebi.ac.uk/Tools/msa/tcoffee/). These alignments were curated manually and the figures were prepared using ESPRIPT server (https://espript.ibcp.fr/ESPript/ESPript/).

## Supporting information

Supplementary_movies_Zipped_folder

## Acknowledgement

PK, NP, IR and VM thank Indian Institute of Technology Bombay (IIT Bombay) for financial support; AD thanks University Grant Commission (UGC), India and ABS thanks Department of Biotechnology, Ministry of Science and Technology, India for their fellowship. The work was supported by the Ramalingaswami Re-entry Fellowship (DBT), a research seed grant from IRCC, IIT Bombay to Prasenjit Bhaumik. We would like to thank Spacetime High Performance Computing (HPC) resource in IIT Bombay, Mumbai, India for generous computing time. The ‘Protein Crystallography Facility’ at IIT Bombay is also acknowledged.This research used resources of the Advanced Photon Source, a U.S. Department of Energy (DOE) Office of Science User Facility operated for the DOE Office of Science by Argonne National Laboratory under Contract No. DE-AC02-06CH11357. Use of the LS-CAT Sector 21 was supported by the Michigan Economic Development Corporation and the Michigan Technology Tri-Corridor (Grant 085P1000817). This project has been funded in whole or in part with Federal funds from the National Institute of Allergy and Infectious Diseases, National Institutes of Health, Department of Health and Human Services, under Contract No.: HHSN272201700059C.

## Author Contributions

Project investigation: DDL, PB; Protein expression and purification: AD, DDL, NP, ABS; Crystal structure solution: JA, TEE; Crystal structure refinement and analysis: JA, TEE, PK, AD, IR, VM, PB; Homology modeling, molecular dynamics (MD) simulation and sequence analysis: PK, AD; Thermal stability assay: ML; Manuscript first draft writing: PK, AD, PB; Data analysis, manuscript writing and reviewing: PK, AD, NP, ABS, IR, VM, JHD, HX, AG, JA, ML, RYY, AW, TEE, DDL, PB.

## Data Availability

X-ray crystallographic coordinates and the structure factor have been deposited in wwPDB with the accession code 7RY7.

## Extended data Figures

**Extended data Figure 1.**
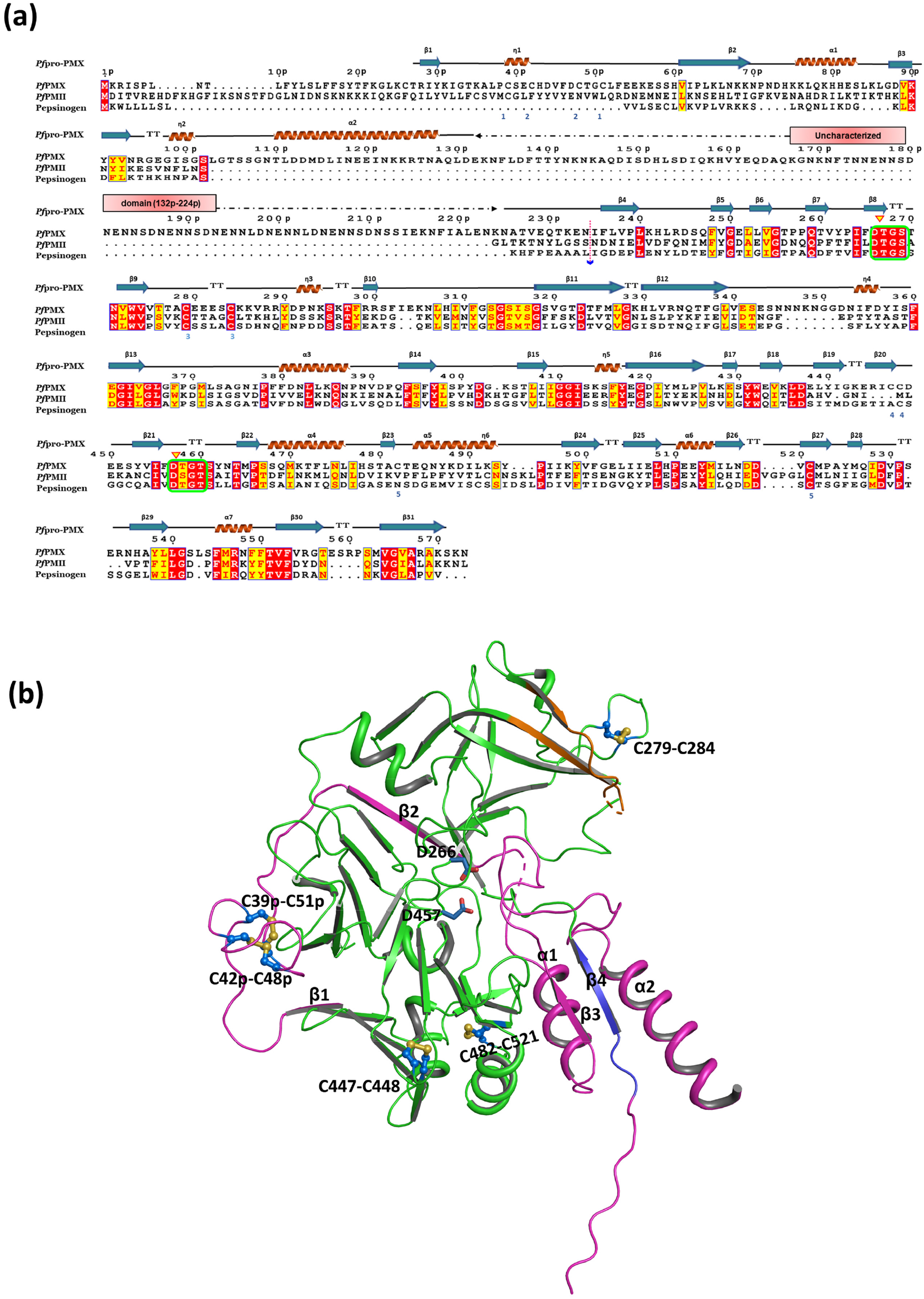
**(a)** Sequence alignment of full-length polypeptide of *Pf*PMX, *Pf*PMII and porcine pepsin zymogens. The secondary structural elements of the *Pf*pro-PMX structure are shown on the top of the sequence alignment. The identical and similar residues are highlighted using ESPript (www.espript.ibcp.fr), identical residues are shown as a white color with a red background, and the similar residues are red characters in yellow background. The numbering of residues of *Pf*pro-PMX is indicated on top and ‘p’ in the numbering refers to the residues from prosegment. The start site of the mature protease is marked by a dotted pink arrow. The active site motif is marked with a green box and catalytic Asp (D266 and D457) with yellow triangles. Blue numbers at the bottom of the alignment indicate the disulfide bonds in the *Pf*pro-PMX structure. The UD is shown with dashed line with pink box on top. (**b**) The disulfide linkages (red stick) in *Pf*pro-PMX structure with the corresponding cysteine residues. The cartoon of the molecule with prosegment in pink, the mature part in green, the flap in orange, and the first β-strand of the mature segment in violet. The secondary structural elements of prosegment are labelled, and catalytic residues (D266 and D457, blue) are shown.

**Extended data Figure 2.**
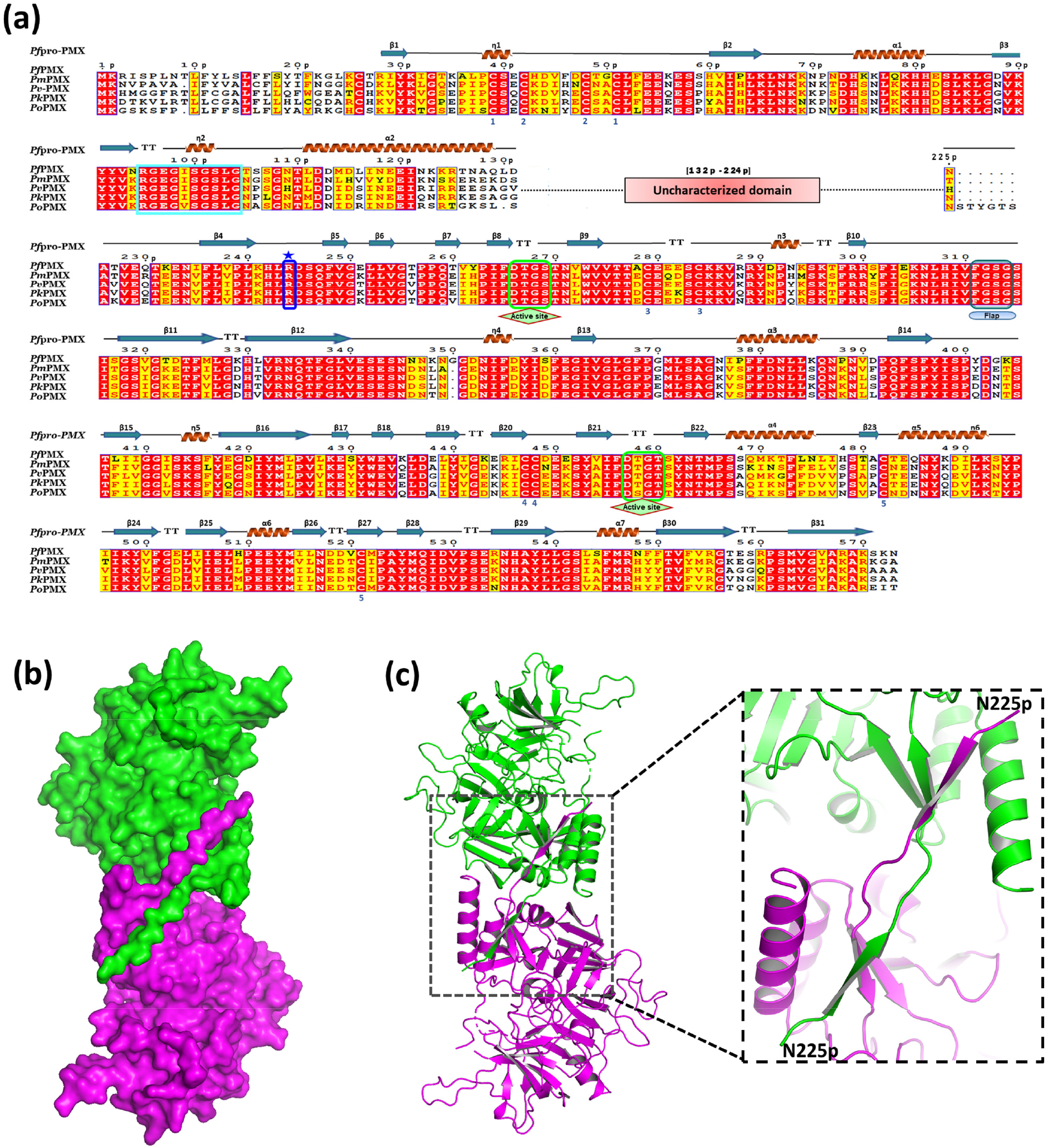
(**a**) Sequence comparison of PMX zymogens from different malaria causing parasites [*P. falciparum* (Pf), *P. vivax* (Pv), *P. knowlesi* (Pk), *P. ovalae* (Po)]. The secondary structure elements of the *Pf*pro-PMX are shown on the top. The identical residues are shown as white characters in a red background, and similar residues are with red characters in the yellow background and disulfide bonds with blue numbers at the bottom of the alignment. The active site motifs are marked with green boxes and flap region with a blue box. The twisted loop region is enclosed within the cyan box and the inhibitory R244 is highlighted with a blue star. (**b**) Surface representation of domain-swapped ‘S’ shaped dimer of *Pf*pro-PMX with one monomer in magenta, and another is in green. (**c**) The cartoon representation of the ‘S’ shaped dimer and the inset shows the swapped loop and the β-strand of the two monomers forming the dimer interface.

**Extended data Figure 3.**
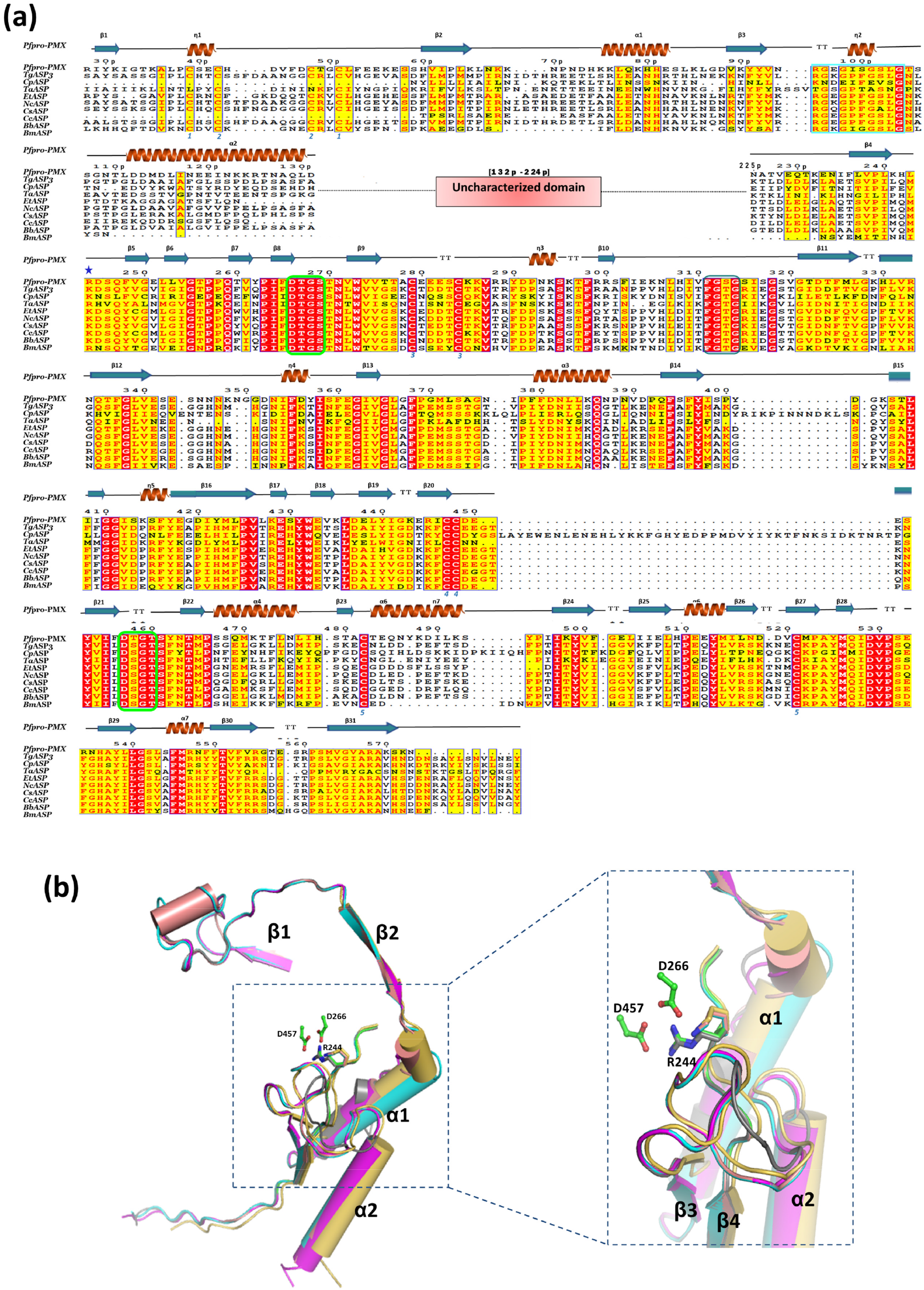
**(a)** Multiple sequence alignment of PMX-like protease zymogens from apicomplexan parasites. The proteases from parasites used in the alignment are: *Plasmodium falciparum* (*Pf*PMX, XP_001349441.1), *Toxoplasma gondii* (*Tg*ASP3, XP_002367043.1), *Cryptosporidium parvum* (*Cp*ASP, XP_627444.1), *Theileria annulata* (*Ta*ASP, XP_954092.1), *Eimeria tenella* (*Et*ASP, XP_013233972.1), *Neospora caninum* (*Nc*ASP, XP_003885934.1), *Cystoisospora suis* (*Cs*ASP, PHJ19229.1), *Cyclospora cayetanensis* (*Cc*ASP, OEH78578.1), *Besnoitia besnoiti* (*Bb*ASP, XP_029222308.1), *Babesia microti* (*Bm*ASP, XP_021337801.1). The identical residues are shown in white with a red background, and similar residues are with red characters in yellow background. Inhibitory arginine /lysis is shown with blue star, twisted loop region is enclosed within the cyan box and catalytic motif is shown with green box. The figure was generated using ESPript. The N-terminal part of the sequences of all the proteases are truncated with respect to R28p of *Pf*pro-PMX. The secondary structural elements of *Pf*pro-PMX are depicted on top of the sequence alignment. Disulfide bonds are shown with blue numbers at the bottom of the alignment, and the uncharacterized domain is shown by the pink box. **(b)** The superposition of the prosegments of *Pf*pro-PMX (magenta prosegment) with the prosegment structures from the models of PMX-like protease zymogens (*Tg*ASP3: cyan; *Cp*ASP: golden; *Ta*ASP; grey and *Et*ASP: salmon) from other parasites. The secondary structural elements of prosegment are labelled. The catalytic aspartates of *Pf*pro-PMX are shown as green colored ball and stick. Inhibitory arginine/lysine are shown as sticks. Inset highlights the similar inactivation mechanism among these proteases through blockage of the active site pocket by twisted loop and salt bridge interactions between arginine/lysine and catalytic aspartates.

**Extended data Figure 4.**
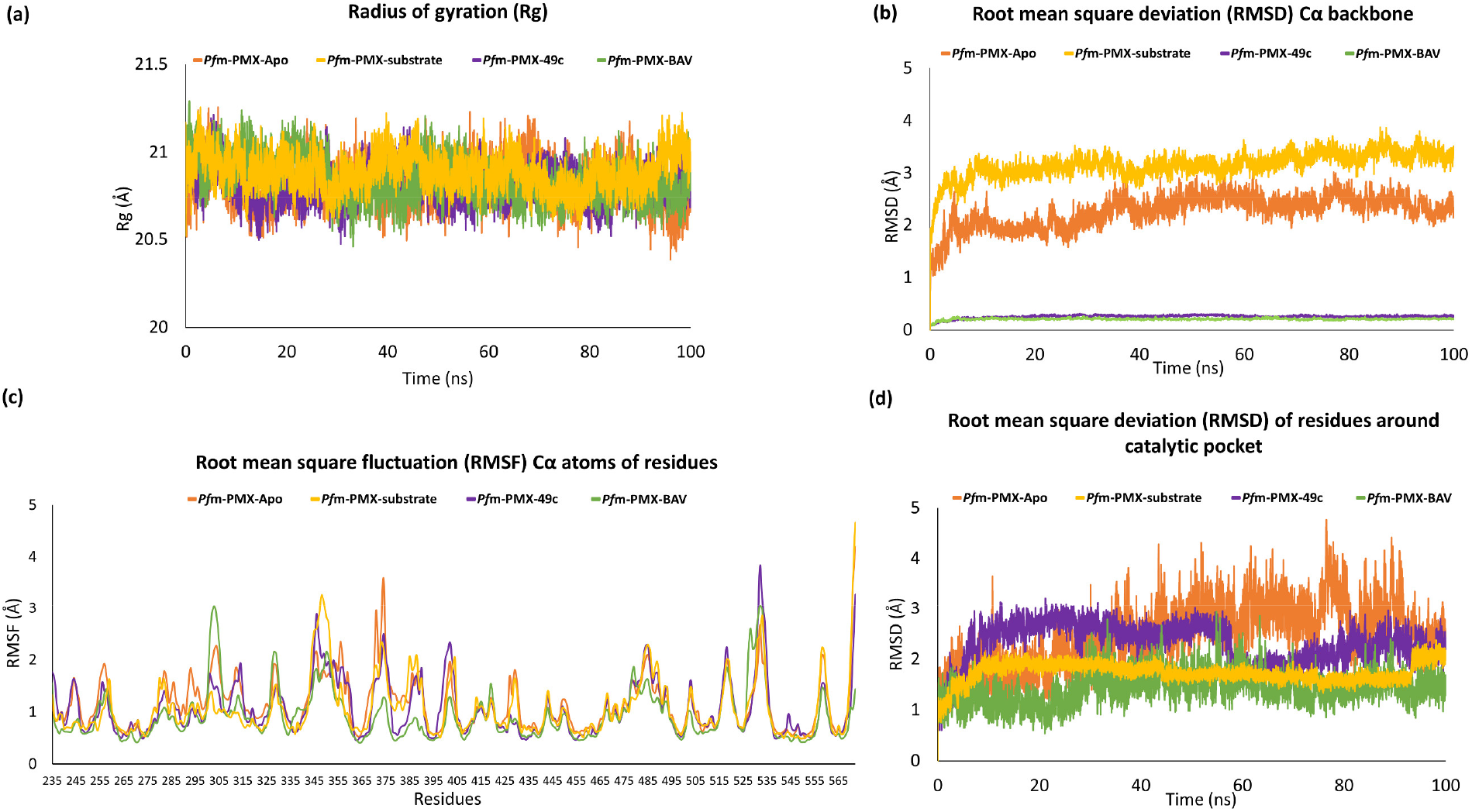
The stability of the overall structures and the active site of *Pf*m-PMX-Apo, *Pf*m-PMX-substrate, *Pf*m-PMX-49c and *Pf*m-PMX-BAV complexes indicated by (**a**) Radius of gyration (Rg) graph, **(b)** Root mean square deviation (RMSD) plot of Cα backbone atoms, (**c**) Root mean square fluctuation (RMSF) of Cα atoms (**d**) The Root mean square deviation plot of the residues around (5.0Å radius) the catalytic aspartates.

**Extended data Figure 5.**
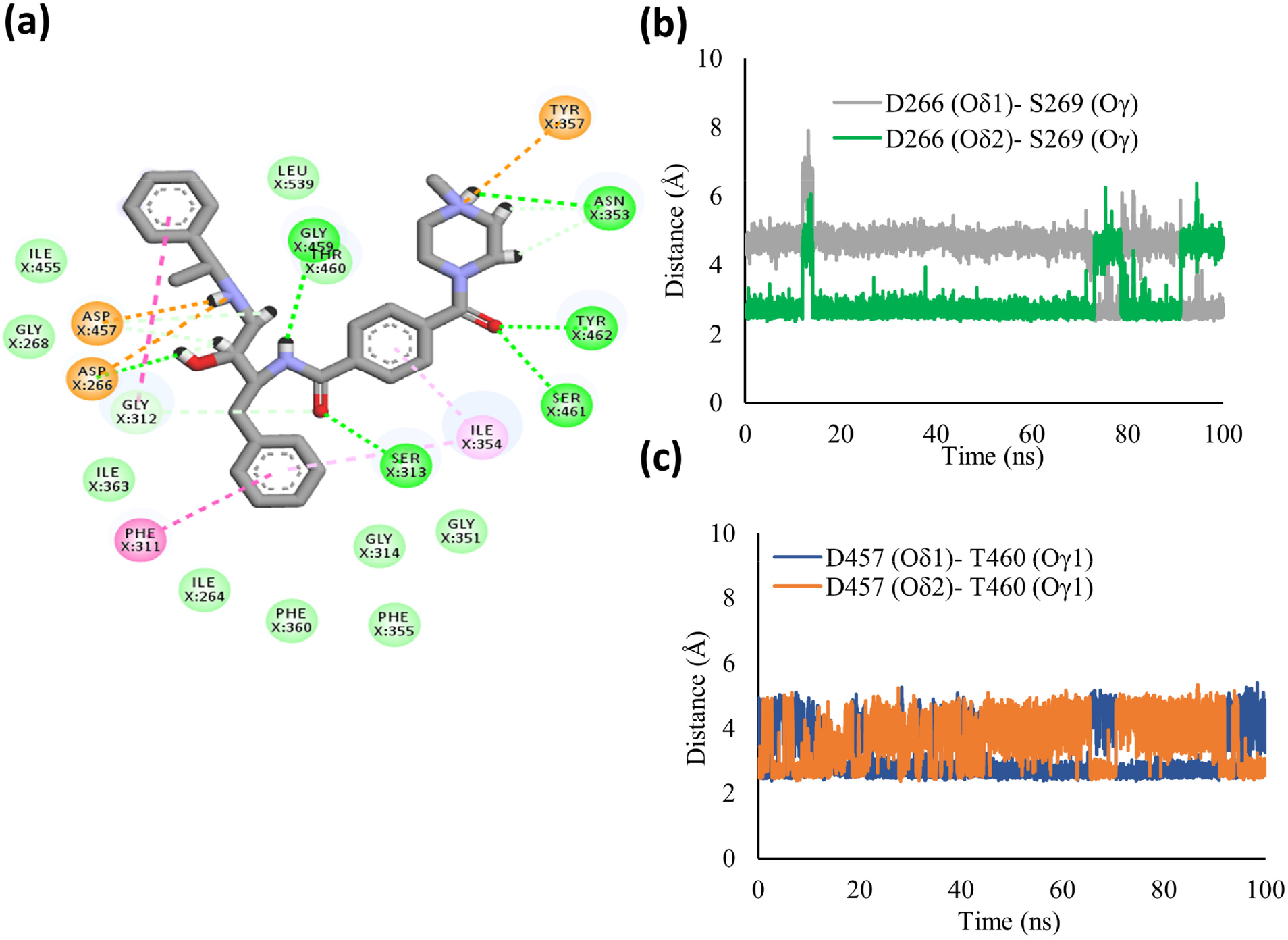
**(a)** The 2D plot of *Pf*m*-*PMX-49c complex showing van der Waals interactions (green sphere), π-π interaction (dark pink sphere), hydrogen bonds (dark green sphere) and salt bridges (orange sphere). The correct orientations of the catalytic aspartates in the *Pf*m*-*PMX-49c complex as monitored by the distances. **(b)** D266(Oδ1-Oδ2)-S269(Oγ) and **(c)** D457(Oδ1-Oδ2)-T460(Oγ1) throughout the simulation.

## Supplementary Materials

### Supplementary movie legends

**Supplementary movie 1**.

*Pf*pro-PMX structure with 2*F*_*o*_*-F*_*c*_ electron density (1σ contour level) of the prosegment. No density is observed for the uncharacterized domain (UD). The mature enzyme portion is presented as surface view and the prosegment is shown as a cartoon in light pink, uncharacterized domain shown with dotted line.

**Supplementary movie 2**

Structural comparison of *Pf*pro-PMX with other pepsin-like aspartic protease zymogens. A movie showing the structural alignment of *Pf*pro-PMX (prosegment in dark nobelium and mature enzyme in light pink) with *Pf*PMII zymogen (prosegment in dark osmium and mature enzyme in light blue) and pepsinogen (prosegment in golden and mature enzyme in light yellow color). The different secondary structural elements of the prosegment and twisted loop (electrostatic surface) of *Pf*pro-PMX are highlighted. Catalytic aspartates are shown as ball and stick. The secondary structural elements of *Pf*pro-PMX prosegment are marked.

**Supplementary movie 3**

The comparison between inhibitory R and K residues in other pepsin-like aspartic proteases. A movie comparing the position of inhibitory Arginine (R244) or Lysine residues (in sticks) between *Pf*pro-PMX (prosegment in dark nobelium and mature enzyme in light pink), phytepsin zymogen (prosegment in dark green and mature enzyme in light green) and pepsinogen (prosegment in golden and mature enzyme in light yellow color).

**Supplementary movie 4**

The movie showing the Rh2N peptide (^P3^HSF/IQ^P2’^) (deep salmon ball andstick) binding in the active site of *Pf*m-PMX. The protein is shown in surface as well as cartoon representation with interacting residues in teal ball and stick and the catalytic aspartates in magenta ball and stick. All the interacting residues have been labelled. Hydrogen bonds are shown by black dashes.

**Supplementary movie 5**

Structural changes during the activation of *Pf*pro-PMX to *Pf*m-PMX. Structure of *Pf*pro-PMX in wheat cartoon and prosegment is in magenta, first β-strand (β1) of mature enzyme (*Pf*m-PMX) is in red color, flap is in orange. Catalytic aspartates, R244 and Y357 are in ball and stick representation and labelled. Uncharacterized domain in magenta dash line.

**Supplementary movie 6**

The binding of 49c (golden ball and stick) in the active site of *Pf*m-PMX. The protein is shown in surface as well as cartoon representation with the interacting residues in teal ball and stick, and catalytic aspartates in magenta ball and stick. All the interacting residues have been labelled. Hydrogen bonds are shown by black dashes.

**Supplementary Figure 1.**
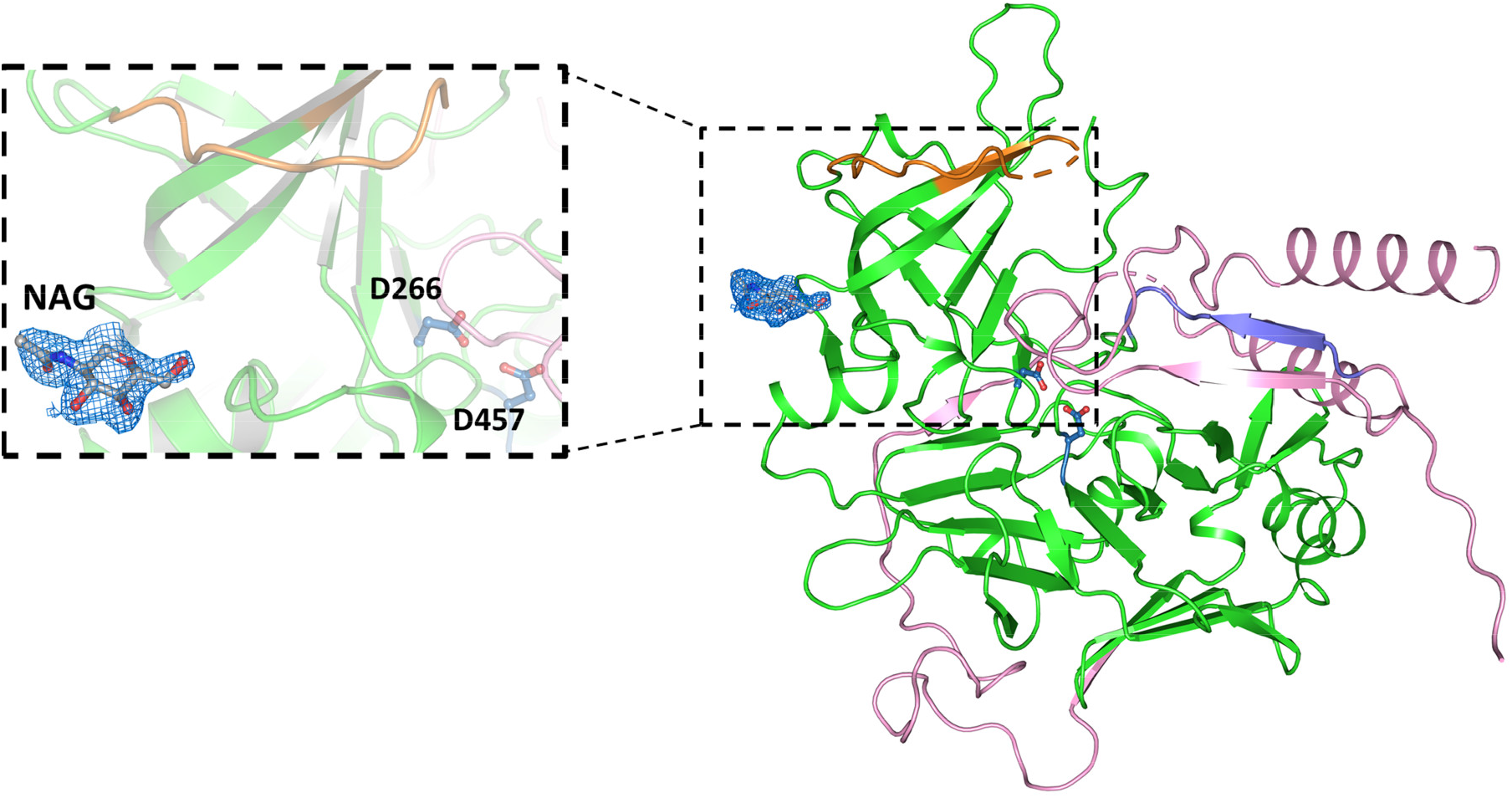
*2F*_*o*_*-F*_*c*_ electron density map (blue mesh) contoured at 1σ level shown around an N-acetylglucosamine (NAG) molecule (grey stick) bound to *Pf*pro-PMX structure. Inset shows the zoomed-in view of bound NAG. The mature enzyme is in green, prosegment is in pink and the flap is orange. The catalytic aspartates are shown in blue sticks.

**Supplementary Figure 2.**
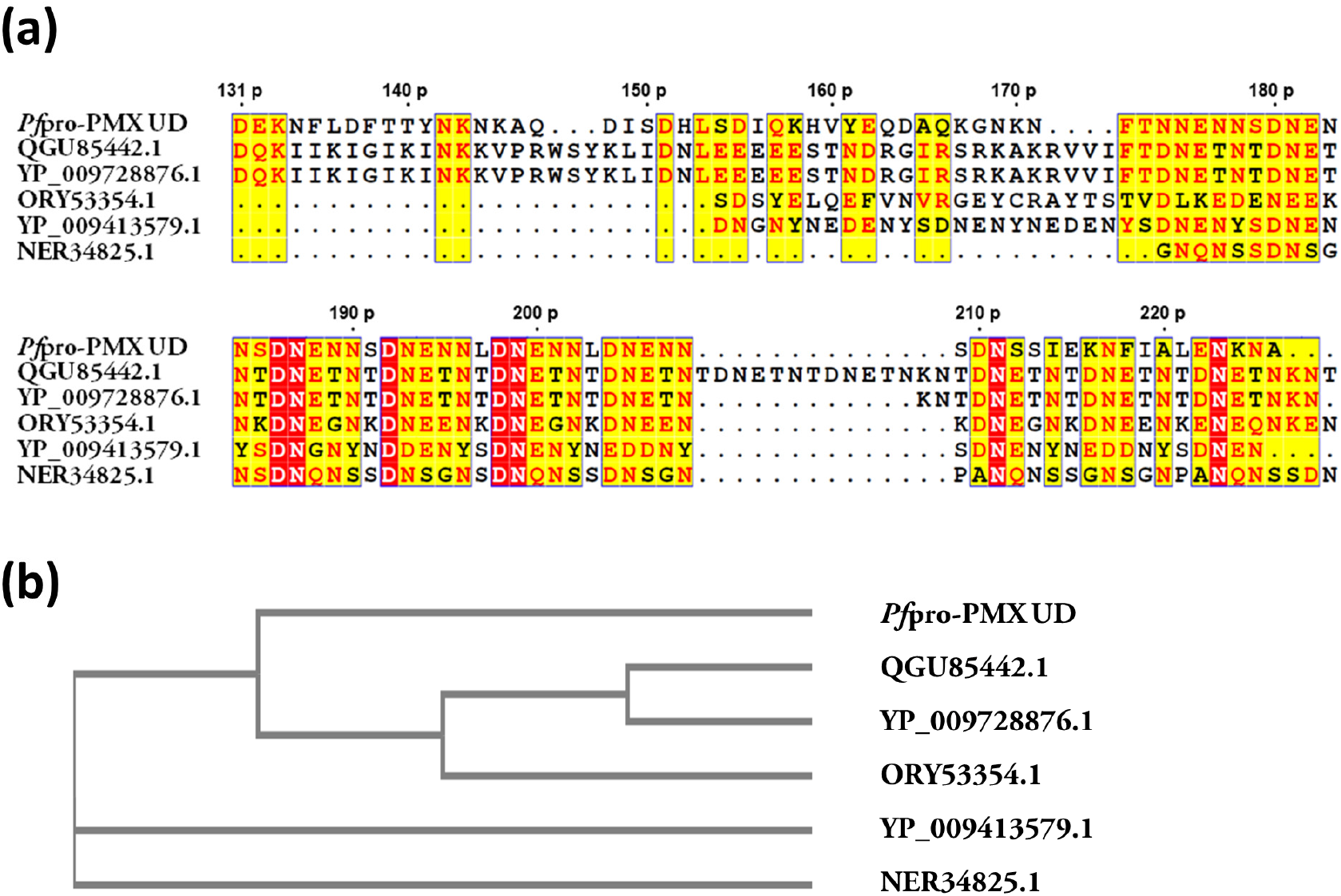
**(a)** Multiple sequence alignment of uncharacterized domain of *Pf*pro-PMX (*Pf*pro-PMX UD) with Photosystem I assembly protein Ycf1 (QGU85442.1) and chloroplast RF1 (YP_009728876.1) from *Anemone cernua*, CHAT domain-containing protein from *Oscillatoria* sp. (NER34825.1), ATP-dependent Clp protease proteolytic subunit (plastid) from *Allotropa virgata* (YP_009413579.1) and coth-domain-containing protein from *Neocallimastix californiae* (ORY53354.1). The residue numbers (‘p’ refers to prosegment) of the *Pf*pro-PMX are shown on the top of the alignment. The cladogram suggests that *Pf*pro-PMX–UD has homology to proteins from plant origin.

**Supplementary Figure 3.**
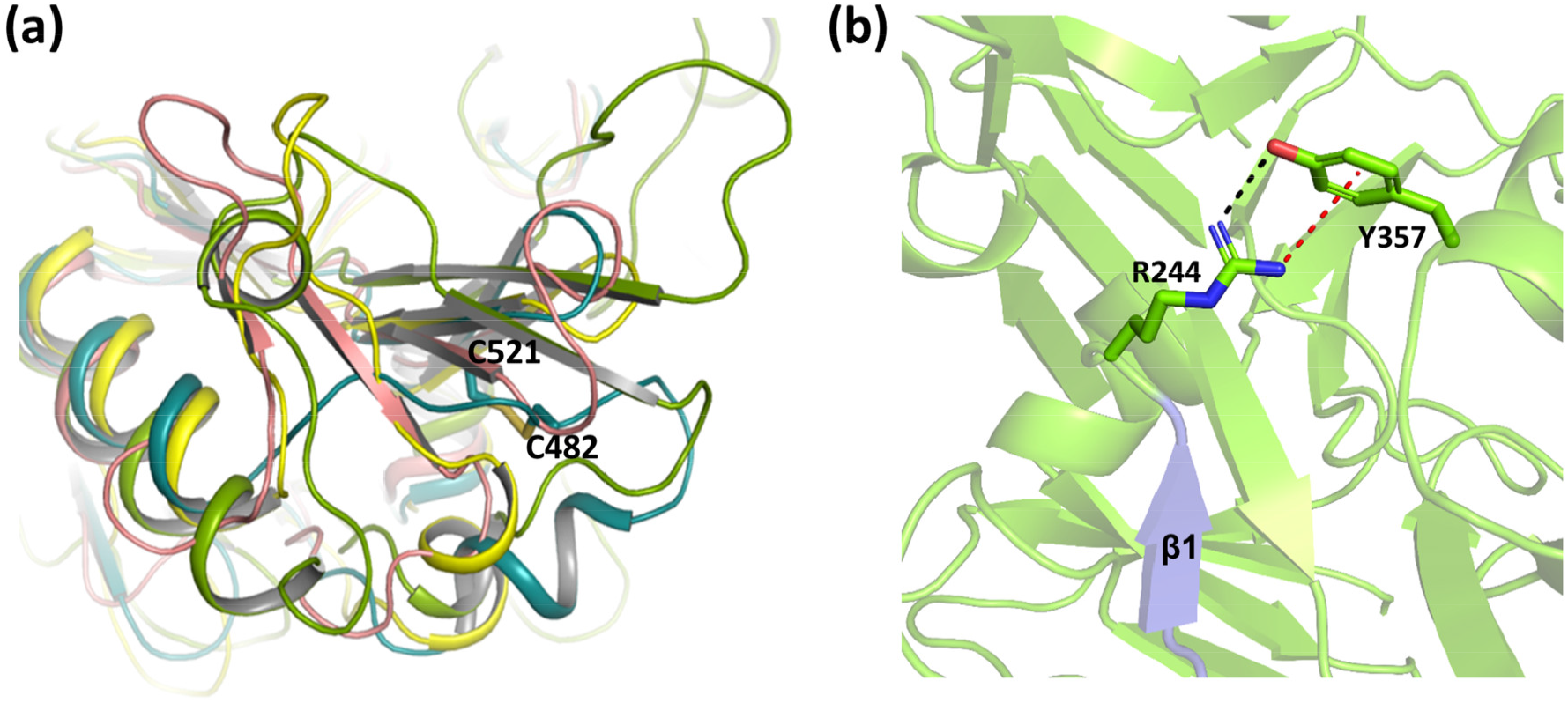
**(a)** Superimposition of zymogen of *Pf*pro-PMX (teal), *Pf*PMII (1XDH, salmon), β-secretase (4GID, green) and pepsin (4PEP, yellow) suggest that the disulphide bond between Cys482-Cys521 which unique to *Pf*pro-PMX maintains the stability of the residues H478-K492 in a short loop and helical conformation. (**b**). In the mature *Pf*m-PMX, R244 forms a cation-π interaction with the side chain of Y357 from the loop N2.

**Supplementary Figure 4.**
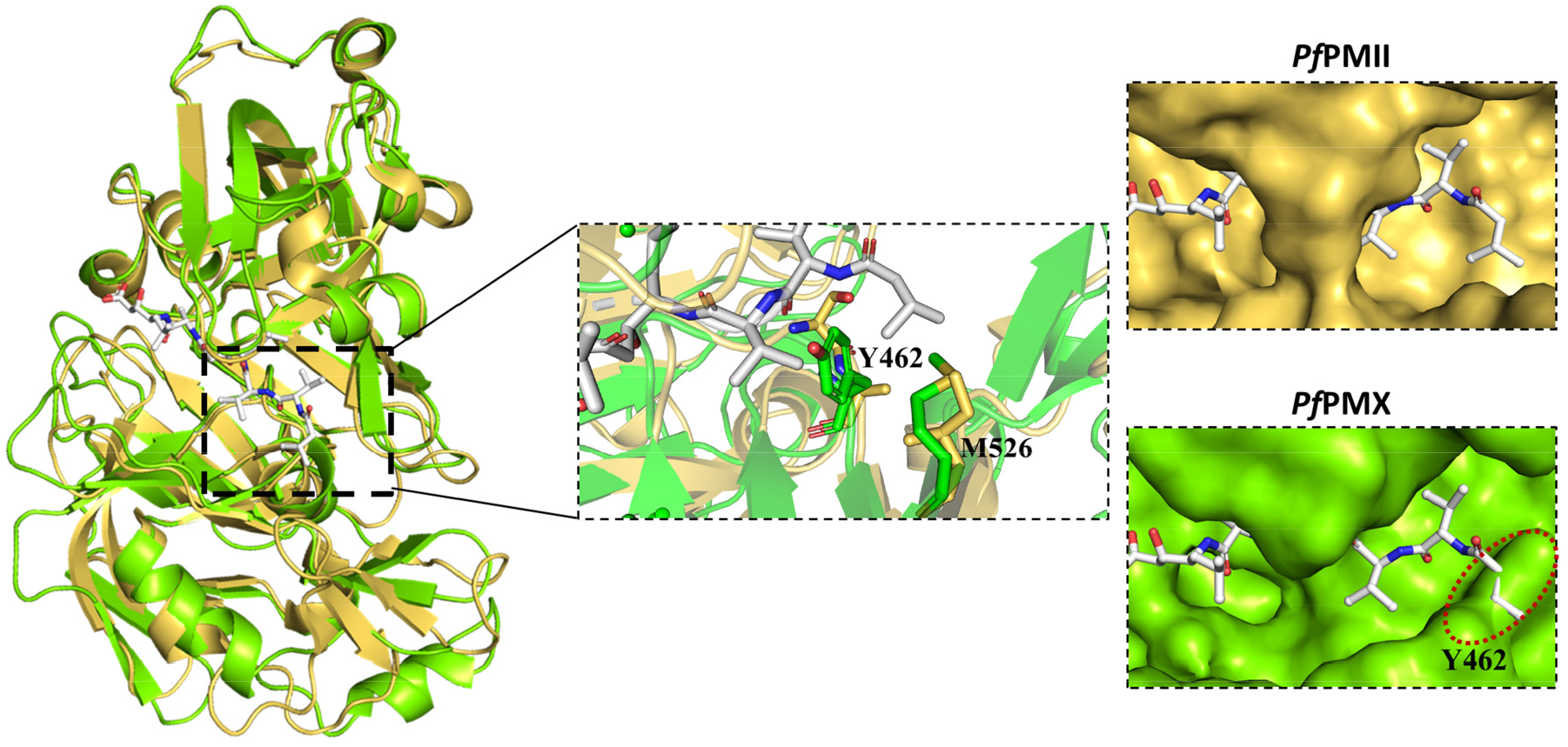
Comparison between the pepstatin A bound complex of *Pf*m-PMX (green) and *Pf*PMII (1M43) (golden). The inset shows the zoomed-in view of the active site indicating Y462 of *Pf*m-PMX occupies the position of A219 of *Pf*PMII and makes the binding pocket narrow. The surface view of *Pf*PMII showing well accommodation of pepstatin A (white stick) in the active site. The surface view of *Pf*m-PMX showing the collision (red dotted circle) of pepstatin A (white stick) with the binding pocket residue Y462 (marked).

### Supplementary data 1

#### Expression and purification of *Pf*PMX in *E. coli*

The optimized gene sequence of *Pf*PMX was synthesized by Thermo Scientific and used for the cloning. Construct 2 (Supplementary Figure 5a) of *Pf*PMX was designed by comparing the protein sequence with *Pf*PMII construct and cloned into pET32b vector between NcoI and Xhol restriction sites containing His_6_ tag for affinity purification, as well as the thioredoxin (TRX) tag to increase protein solubility. The recombinant plasmid of *Pf*PMX was transformed into an expression strain of E. coli SHuffle T7 Express lysY. A single colony expressing a high amount of the recombinant fusion protein was inoculated into 5 ml of LB with ampicillin (100 μg/ml) as an antibiotic selection marker. The culture was grown at 30°C, with shaking at 150 rpm overnight. 1% of the overnight grown culture was used as inoculum for secondary culture with the same antibiotic concentration in LB and grown at 30°C in shaking condition until the OD600 reached 0.6-0.8. The culture was then induced with 0.4 mM isopropyl β-D-1-thiogalactopyranoside (IPTG) and incubated overnight at 16°C for the overexpression of the recombinant protein. Cells were harvested by centrifugation at 8000 rpm for 10 min. The cell pellet was resuspended into the lysis buffer (buffer A: 50 mM Tris, 300 mM NaCl, 10 mM imidazole, 0.2% CHAPS, pH 7.4) and lysed with sonication for 15 min. The lysate was centrifuged at 16,000 g for 30 min and the supernatant containing a soluble fraction of protein was used for purification. The supernatant was then loaded onto a His-Trap affinity column (GE Healthcare) equilibrated with buffer B (50 mM tris, 300 mM NaCl, 0.2%.

Purification of expressed protein from construct 2 shows the presence of full-length fusion protein at ∼61.2 kDa (Supplementary Figure 5b, lane2) along with some impurities. The cleavage of the tags with enterokinase, also results in conversion of fusion protein to a polypeptide of ∼43.3 kDa (Supplementary Figure 5b, lane 3), which, on storage at 4 °C, leads to its conversion to ∼40 kDa form (Supplementary Fig. 5b, lane 4). These observations from both constructs suggests that the prosegment of *Pf*PMX zymogen has at least two cleavage/maturation sites: (a) cleavage site 1 (CS-1) at the start of the UD and (b) cleavage site 2 (CS-2) at the end of the UD. For this construct, we assume that after removal of the tag ∼43.3 kDa zymogen gets cleaved at CS-2 and converts to ∼40 kDa mature domain (Supplementary Fig. 5a).

**Supplementary Figure 5.**
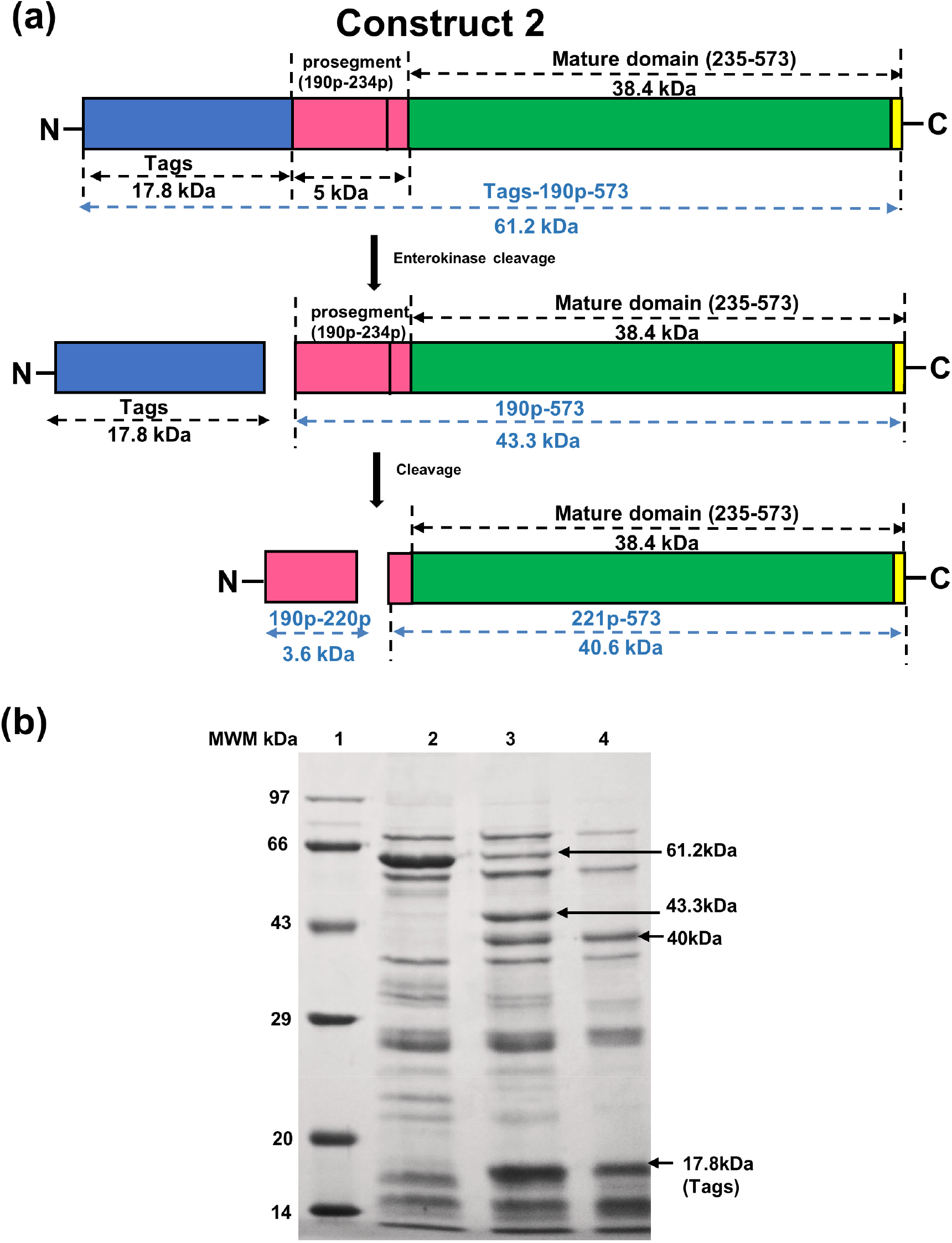
**(a)** Schematic diagram of construct 2 of *Pf*PMX consisting of tags (17.8 kDa includes thioredoxin tag, His_6_ tag, S-tag, and enterokinase cleavage site), prosegment (5 kDa), and mature domain (40 kDa). The removal of tags by enterokinase would form a peptide which upon subsequent cleavage leads to a formation of the mature enzyme. **(b)** SDS-PAGE analysis for eluted fractions of construct 2 after size exclusion chromatography, lane1: molecular weight marker (MWM), lane 2: full length protein, lane 3: zymogen and mature form lane 4: mature protein.

### Supplementary data 2

#### Virtual screening of inhibitors against *Pf*m-PMX

The molecule 49c was docked into the active site of *Pf*m-PMX and the complex was simulated for 100 ns. The interactions of 49c were analysed to understand the structural basis of inhibitor binding in the active site of *Pf*m-PMX. Based on these interactions further screening of multiple inhibitors was performed.

A total of 57 inhibitor molecules were considered. Some of them have been reported to bind to the pepsin-like aspartic proteases and the rest are chemically similar molecules downloaded from the PDB. The inhibitors were docked into the active site of *Pf*m-PMX using AutoDock 4.2^1^. The correct orientations of the inhibitors in the active site were analyzed based on the distance between the hydroxyl group of inhibitors and the catalytic aspartates of *Pf*m-PMX. The docking score corresponding to those selected poses were used for ranking the binding affinities of these molecules (**Supplementary Table S1**). Among the screened inhibitors, β-secretase (BACE-I) inhibitors (BAV and 74A) and HIV-1 protease inhibitor indinavir (MK1) showed the highest binding affinity to *Pf*m-PMX. Most of the BACE-I inhibitors showed good binding affinity towards *Pf*m-PMX. Certain HIV-1 protease inhibitors and some of the KNI compounds also showed comparable binding affinities.

BAV is a macrocyclic peptidomimetic inhibitor of BACE-I, a pepsin-like aspartic protease^2^. The mode of binding of BAV in the active site of *Pf*m-PMX is shown in **Supplementary Figure 6a**. All the distances involving active site residues are similar to the corresponding distances in the apo-*Pf*m-PMX. The BAV in *Pf*m-PMX is oriented in a similar fashion as in the BACE1-BAV crystal structure (3DV5) (**Supplementary Figure 6b**), wherein the central hydroxyl group (O46) makes hydrogen bonds with the carboxyl side chain of D266, and its nearby amine group (N51) interacts with the carboxyl side chain of D457 (**Supplementary Figure 6c and 6d**). The amide group at position N5 interacts with the carbonyl group of the backbone of G459. The macrocycle occupies the S1 pocket and makes hydrophobic contact with the F311 of the flap and the isopropyl benzene group at the other end occupies the S2’ subsite and interacts with Y431 (**Supplementary Figure 6e**). The residues involved in interaction with BAV are conserved among PMX-like proteases, and thus BAV would be a good candidate for drug discovery against these proteases.

**Supplementary Figure 6.**
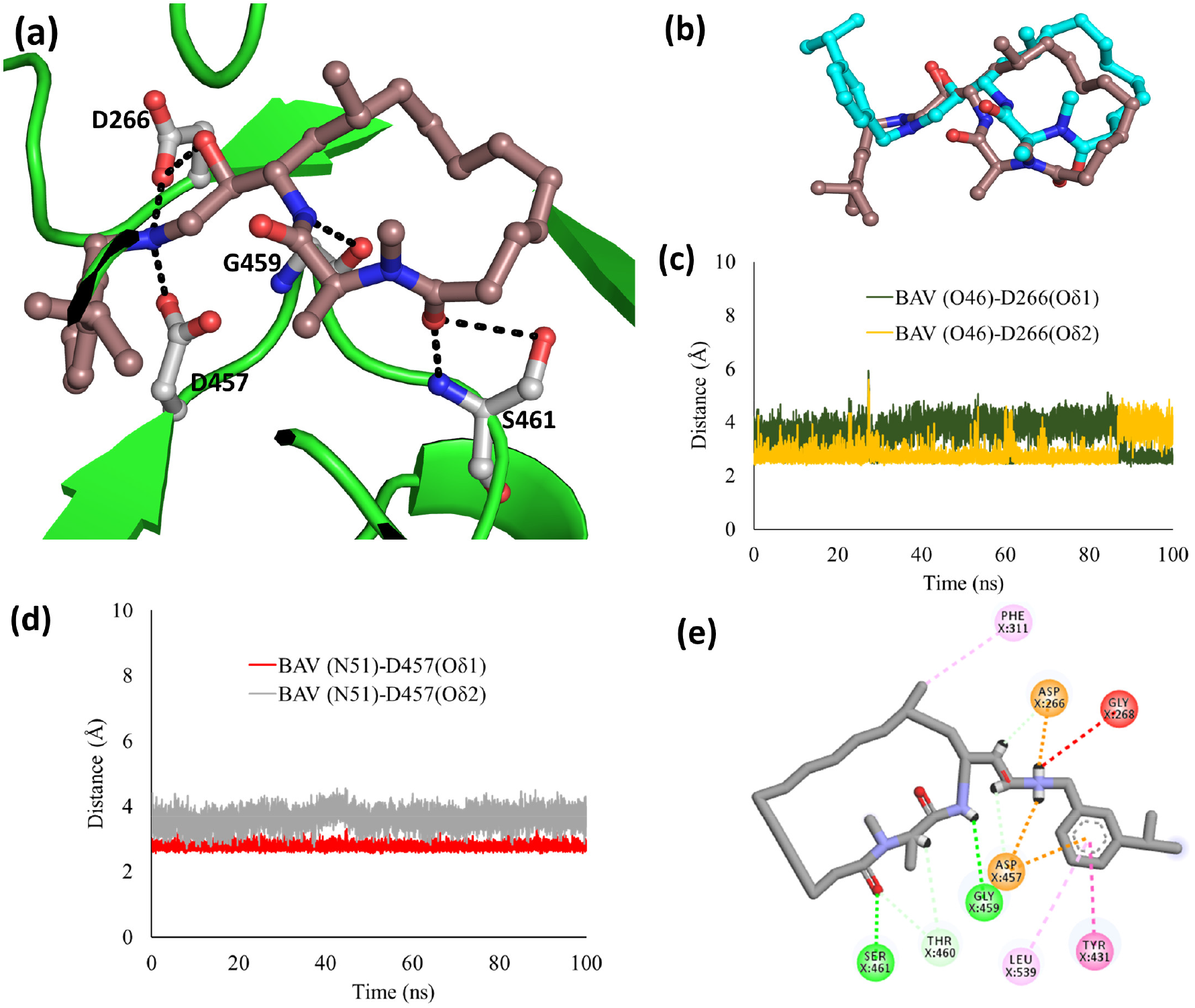
(**a**) Binding mode of BAV (brownish pink) into the active site of *Pf*m-PMX, with residues shown by grey carbon and interactions with black dashes. (**b**) 3D orientation of the BAV in *Pf*m-PMX-BAV complex (brownish pink) as compared to the same inhibitor as observed in the crystal structure of BACE1-BAV (3DV5) (cyan). The block of the catalytic aspartates with the inhibitor in *Pf*m-PMX-BAV complex as indicated by the distances (**c**) BAV(O46)-D266 (Oδ1-Oδ2) and (**d**) BAV(N51)-D457 (Oδ1-Oδ2) throughout the simulation. (**e**) The 2D plot of *Pf*m-PMX-BAV complex showing van der Waals interactions (green sphere), π-π interaction (dark pink sphere), hydrogen bonds (dark green sphere and salt bridges (orange sphere).

**Supplementary Table 1:**
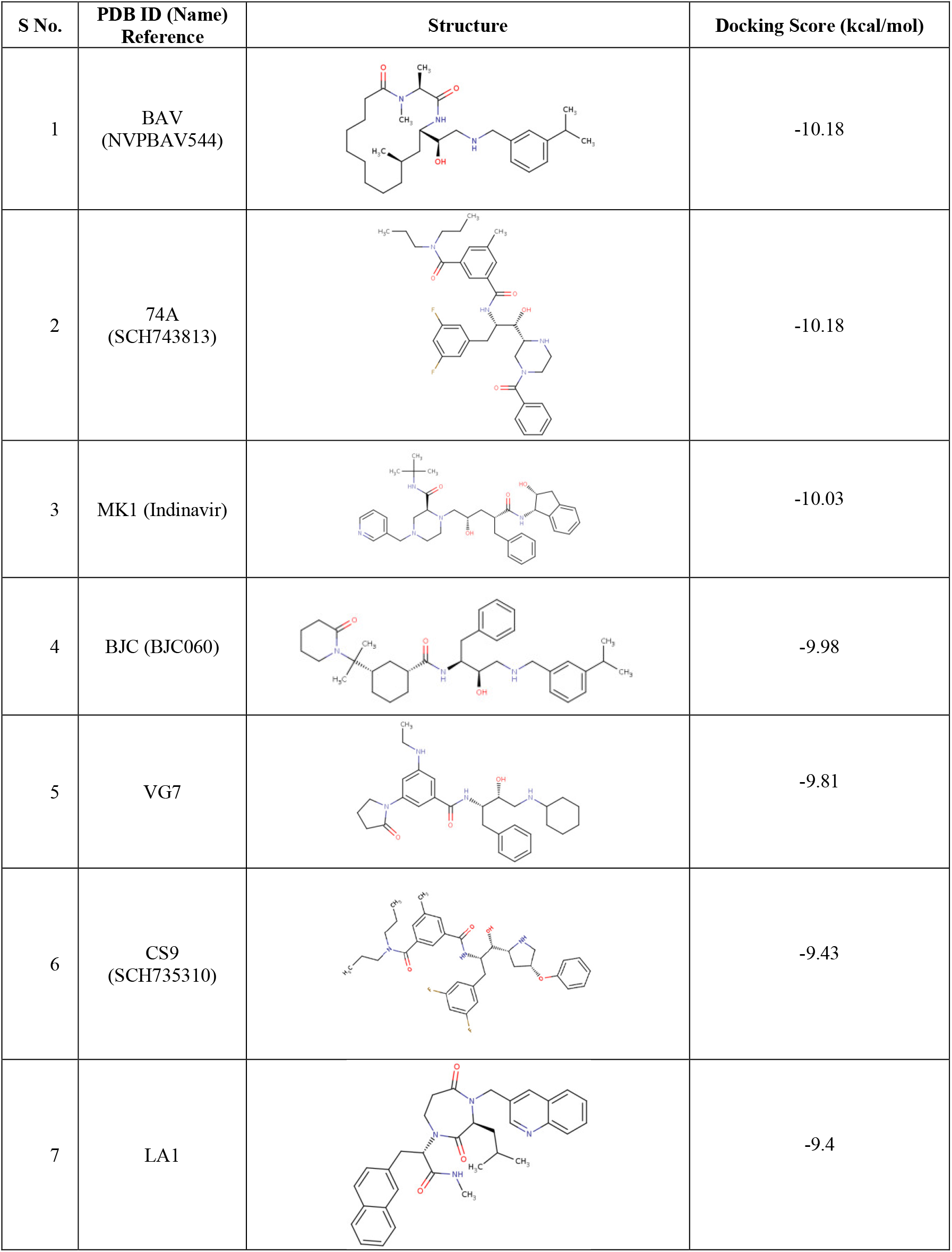
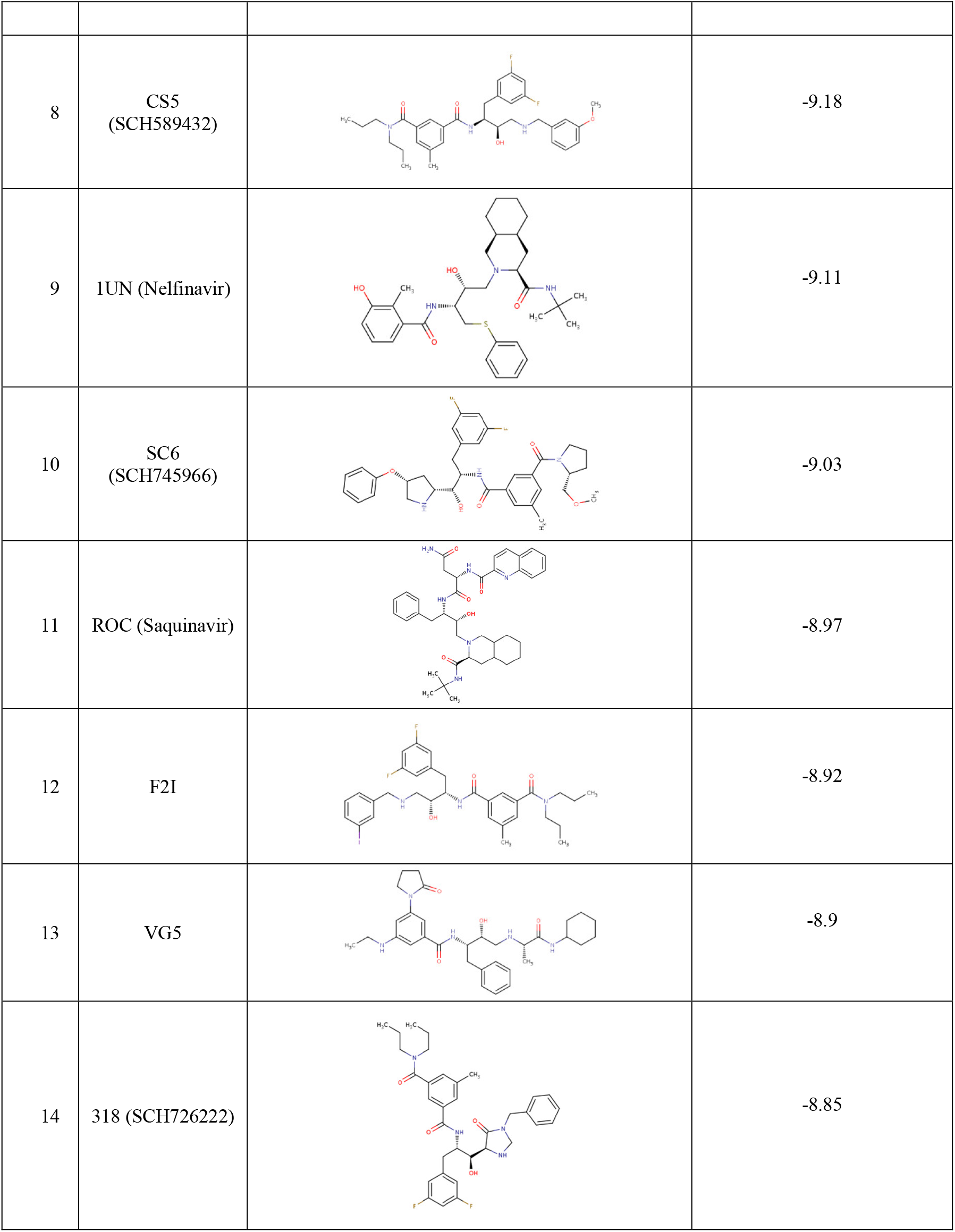
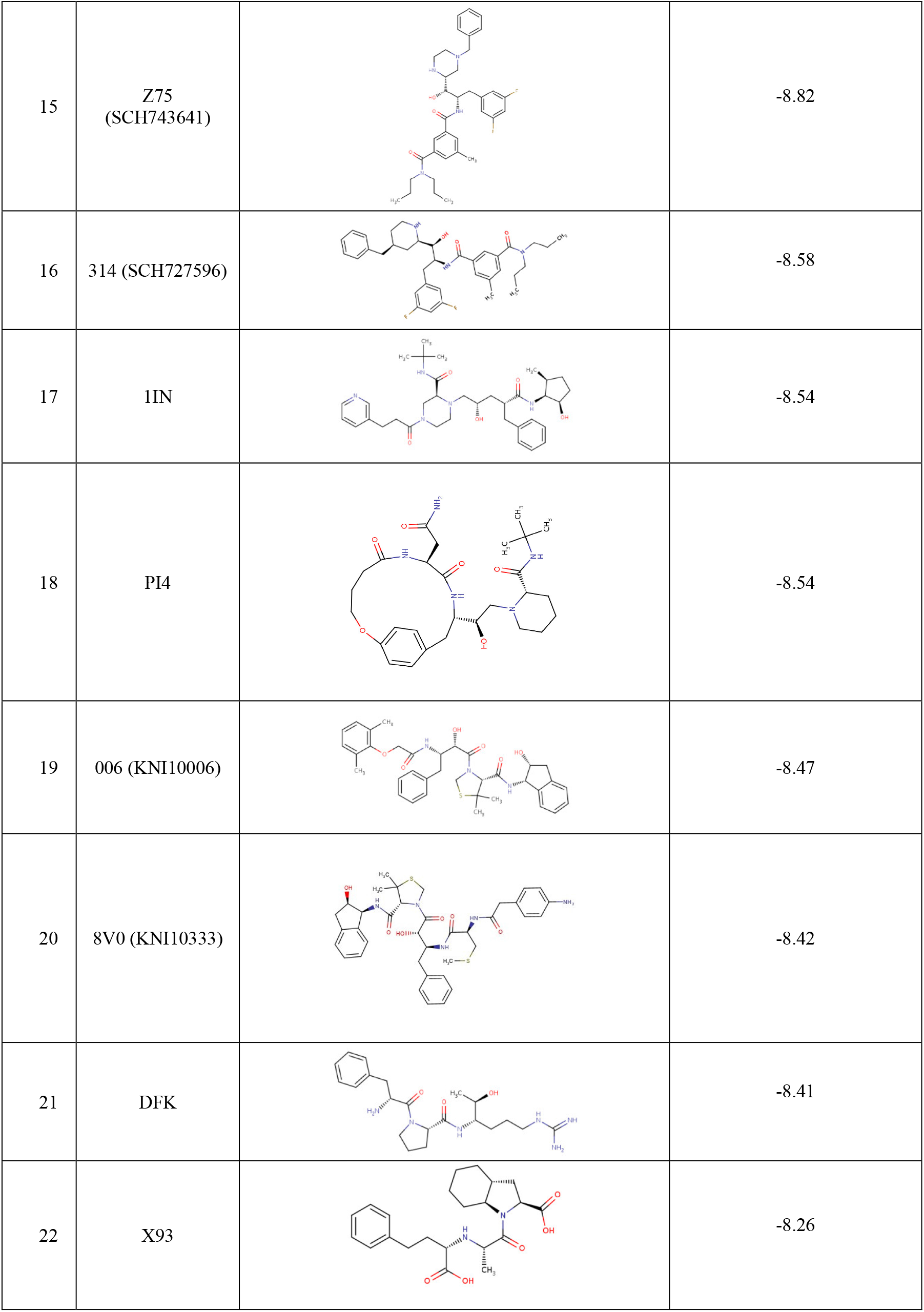
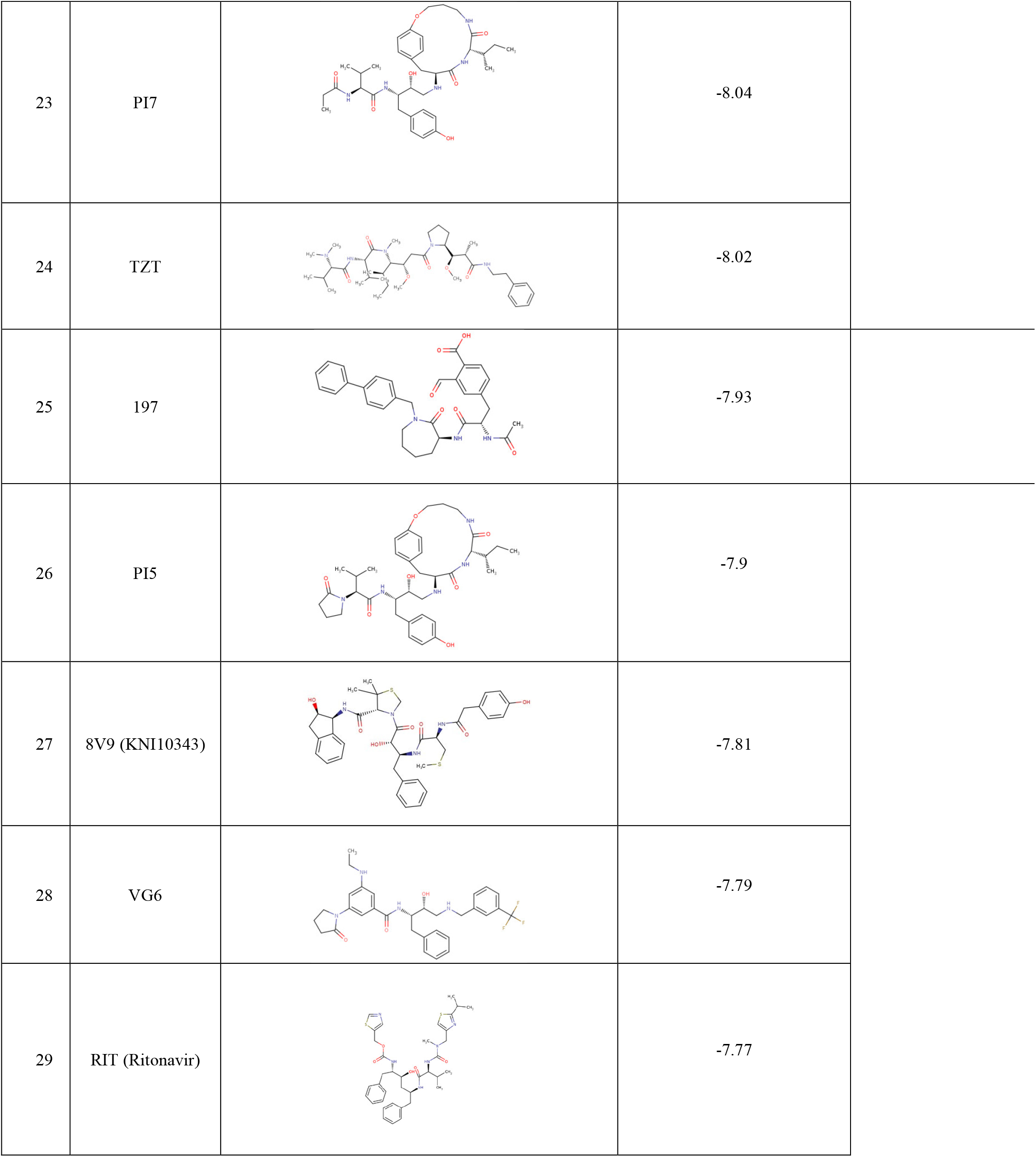
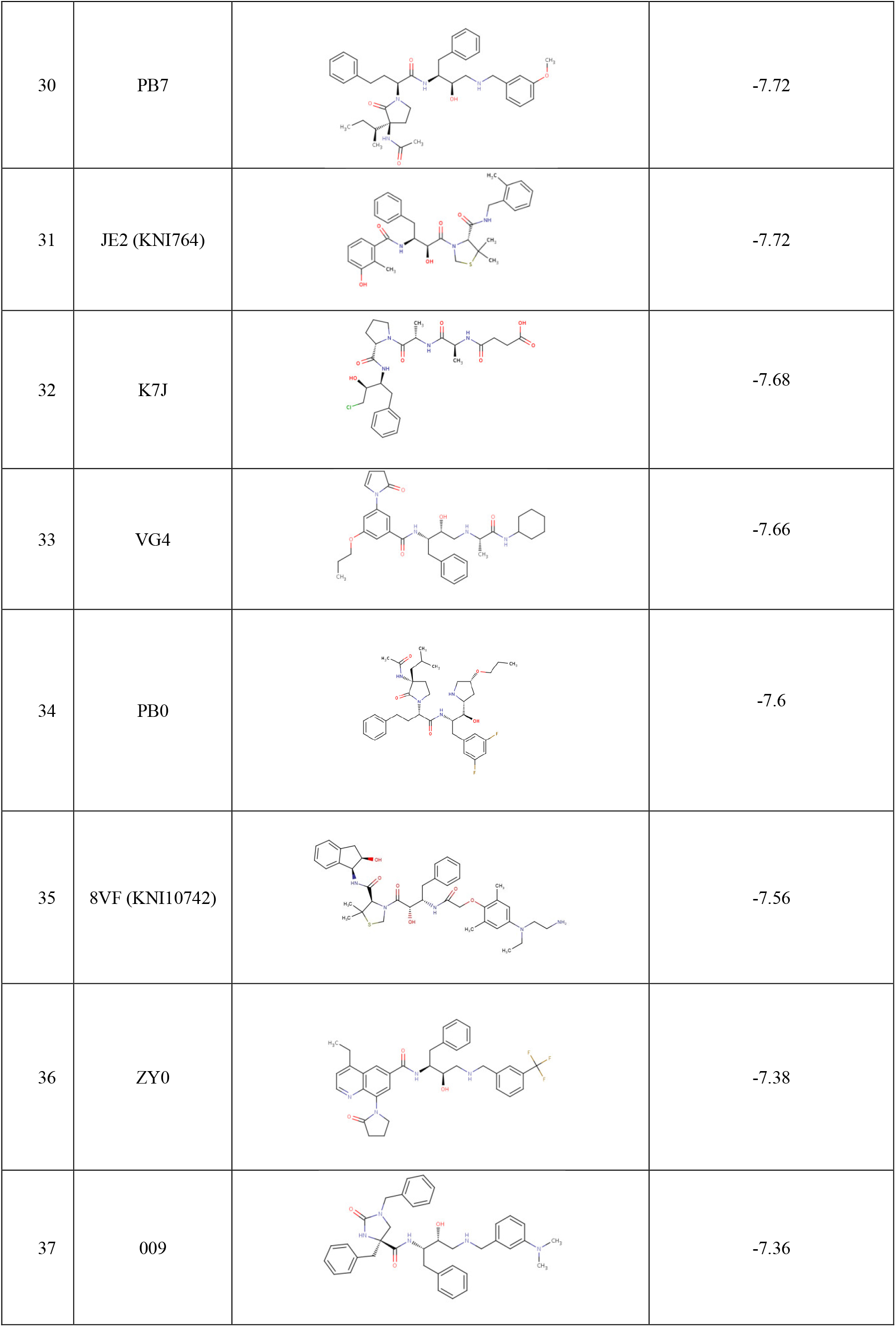
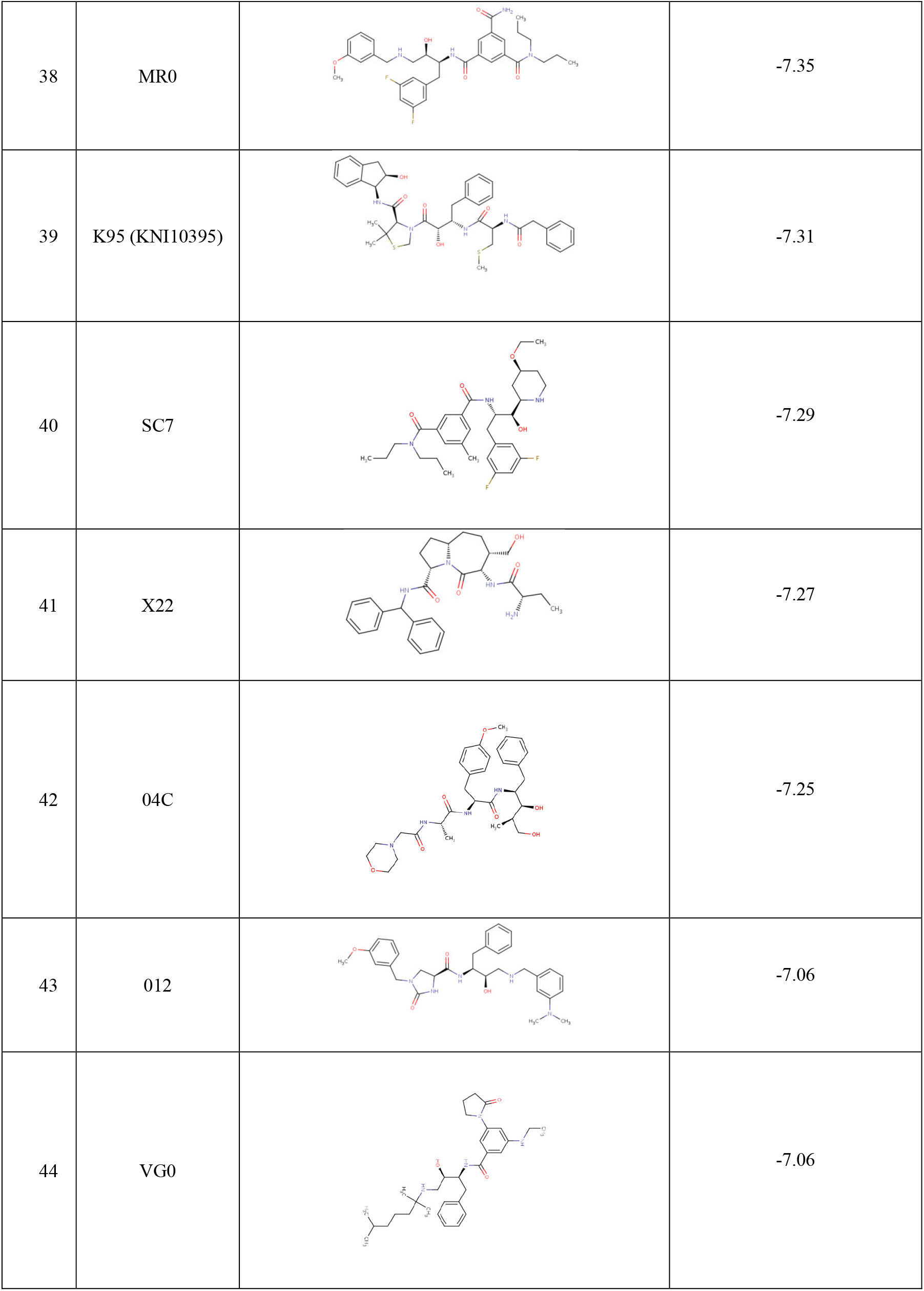
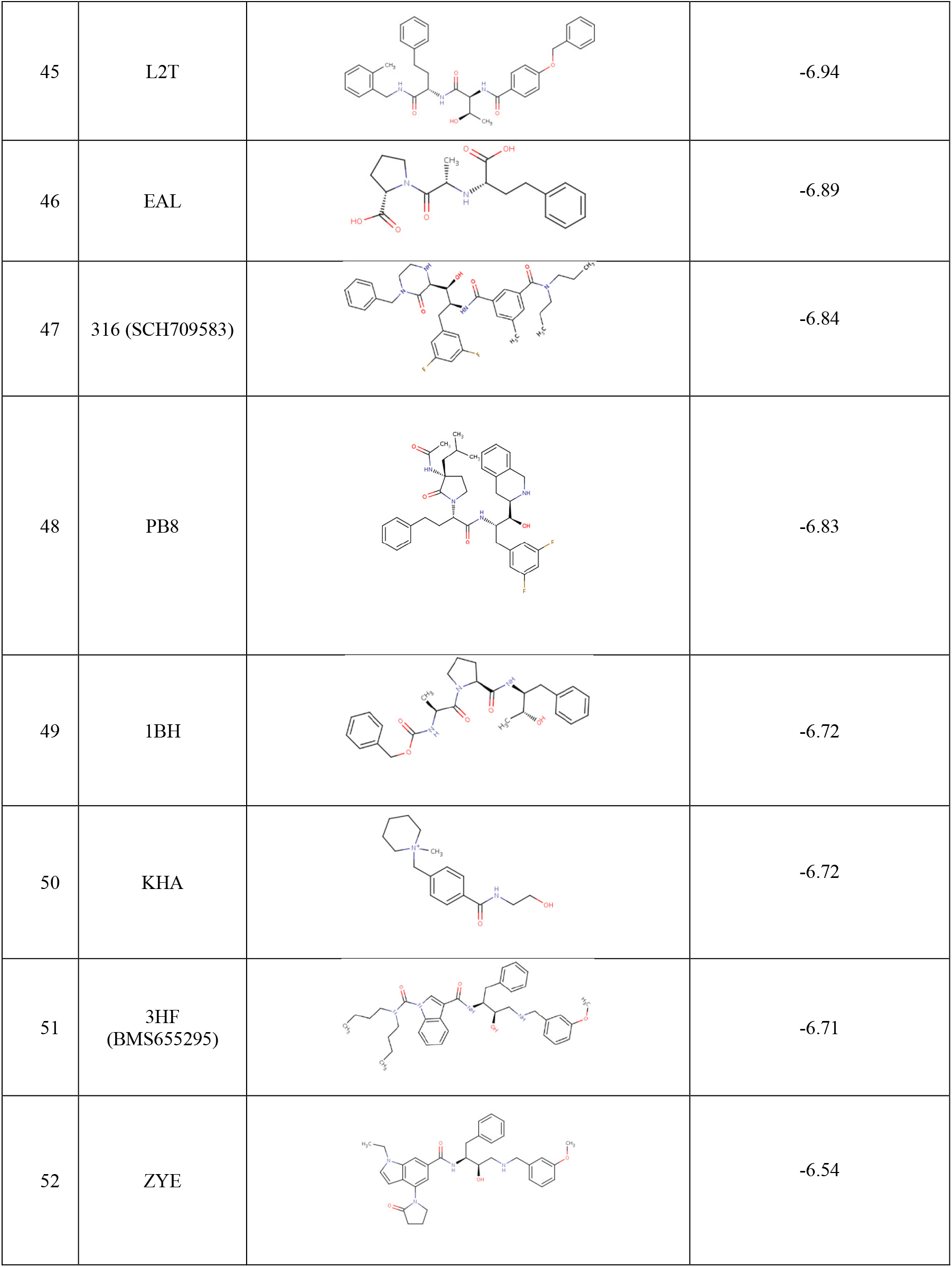
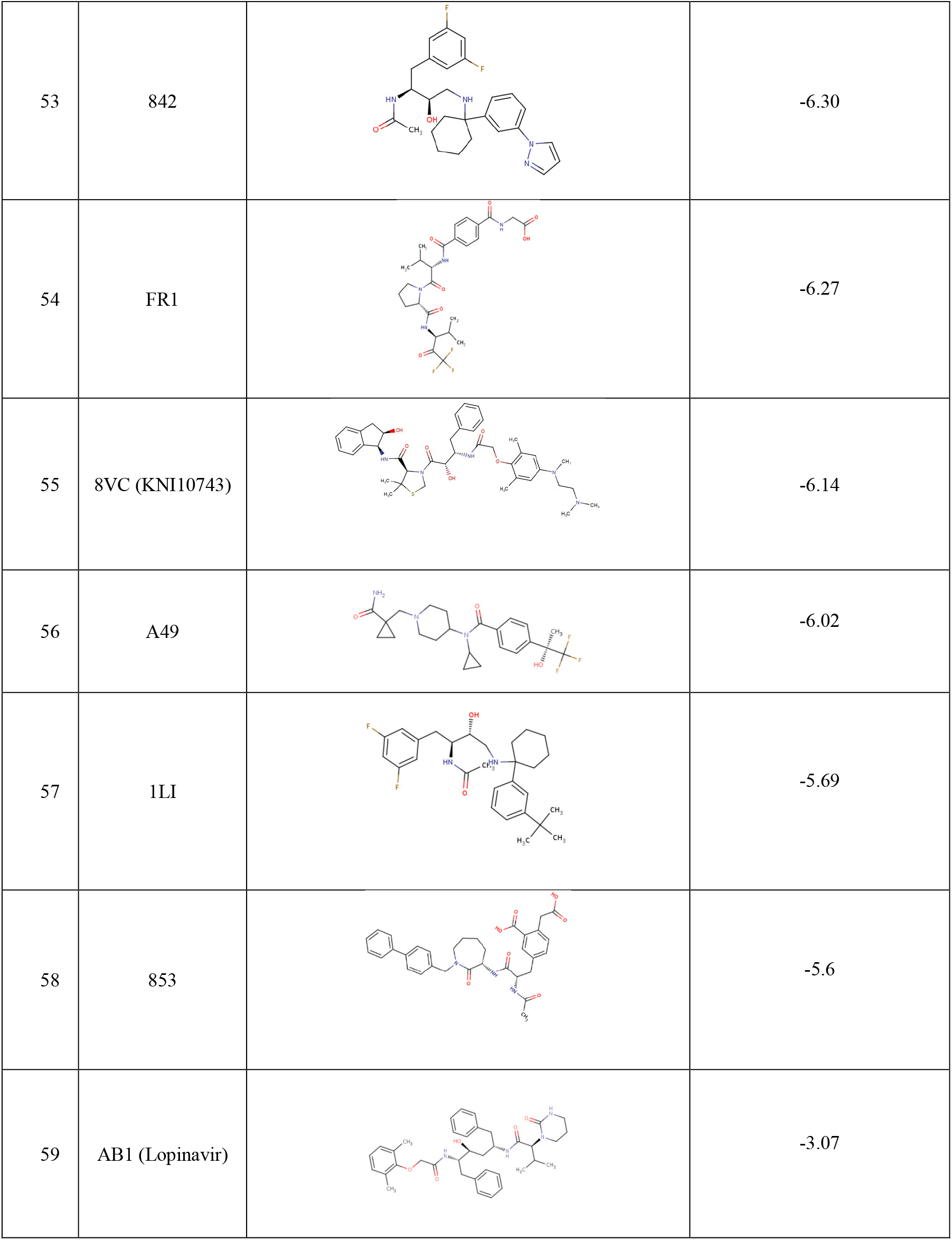
The docking score of the inhibitors screened against *Pf*m-PMX. The molecules are ranked as per docking scores. The covalent structures of the inhibitors and their PDB IDs has also been provided

